# Targeting insulo-frontal pathway to reduce stress-evoked cognitive rigidity

**DOI:** 10.64898/2026.02.12.705479

**Authors:** Shaorong Ma, Kuan Hong Wang, Yi Zuo

## Abstract

The anterior insular cortex (aIC), a central hub of the salience network, is engaged by cognitive flexibility tasks^1,2^; it is also implicated in stress-related mental disorders^3–5^, where cognitive rigidity is a common but poorly treated symptom^1^. While the insular cortex’s roles in interoception and emotional regulation are extensively studied^6–10^, its causal contribution to cognitive rigidity remains unclear. Using attentional set-shifting tasks (AST) in mice, we identify aIC neurons projecting to the medial prefrontal cortex (mPFC) as key regulators of adaptive decision-making. These neurons show heightened activity following incorrect—but not correct—trials. This elevated activity persists into subsequent trials, providing a salience signal that enhances mPFC outcome-dependent updating and promotes convergence of neural activity patterns across trials. Optogenetic manipulation of aIC→mPFC projections during the pre-decision phase disrupts mPFC updating and impairs AST performance. Moreover, stress disrupts the outcome-dependence of aIC activity and impairs set-shifting. Crucially, selectively reinforcing aIC→mPFC activity after incorrect trials via optogenetics enhances mPFC updating, improves neural activity convergence across trials, and restores cognitive flexibility in stressed mice. These findings reveal a previously unrecognized role of the aIC→mPFC circuit in linking trial outcomes to adaptive decision-making and identify this pathway as a promising target for treating stress-induced cognitive rigidity.

## Main

Cognitive flexibility, the ability to adapt mental strategies and behaviors in response to changing rules or information, is a core component of executive functions that underpins effective decision-making and problem-solving in complex settings^11,12^. In today’s fast-paced, dynamic world, such adaptability is critical for managing the escalating demands of decision-making. However, stress—a pervasive feature of modern life—undermines cognitive flexibility^13–15^, promoting rigidity that exacerbates daily challenges and increasing the risk of mental disorders^16,17^. Addressing this issue requires a deeper understanding of how stress impairs cognitive flexibility at the neural circuit level, thereby paving the way for targeted interventions to restore adaptive behavior.

The anterior insular cortex (aIC) serves as a central hub of the salience network (SN), a brain network responsible for detecting and prioritizing salient stimuli and facilitating the dynamic switching between the default mode network (DMN) and the central executive network (CEN)^2,18,19^. Both of these processes are involved in cognitive flexibility^12^. Stress has been shown to cause aIC dysfunction^20,21^, which is associated with deficits in cognitive flexibility^22^ and stress-related neuropsychiatric disorders^23,24^. Furthermore, detailed anatomical studies in non-human animals demonstrate that the aIC directly projects to the medial prefrontal cortex (mPFC)^25,26^, a region critical for decision-making and cognitive flexibility in both humans and animals^27–34^. Yet, it remains unclear how the aIC interacts with the mPFC to regulate cognitive flexibility, and how stress-induced dysfunction of this pathway contributes to cognitive rigidity. Elucidating these circuit-level mechanisms is crucial for understanding the neural basis of stress-related impairments in cognitive flexibility and for developing targeted interventions for individuals affected by stress-induced cognitive deficits.

## Results

### Manipulating aIC→mPFC impairs flexibility

Cognitive flexibility is assessed in humans and animals commonly through attentional set-shifting tasks (AST)^35^, in which subjects are presented with compound sensory cues, and the cue-reward contingency shifts between different sensory features or modalities. A typical rodent AST test consists of a series of sessions: 1) simple discrimination (SD), in which the mouse chooses between two digging media associated with either distinct textures or odors (first sensory modality), with only one texture or odor being associated with food reward; 2) compound discrimination (CD), in which a second sensory modality (*i.e.*, odor is added if the first modality is texture, and *vice versa*) is introduced, but the rewarded cue is the same as in SD; 3) reversal (Rev), which preserves the task-relevant modality but swaps the cues for reward; 4) intra-dimensional shift (IDS), which preserves the task-relevant modality, but replaces the two cues in each modality with novel cues; 5) extra-dimensional shift (EDS), which swaps the former task-relevant and task-irrelevant modalities with all cues replaced. Thus, the animal learns to associate rewards with sensory cues changing within a modality in earlier sessions (SD, CD, Rev, and IDS), but in EDS it needs to shift its attention and decision-making strategy significantly as the cue-reward contingency switches to a different sensory modality (Table 1, Fig. 1a).

**Figure 1.**
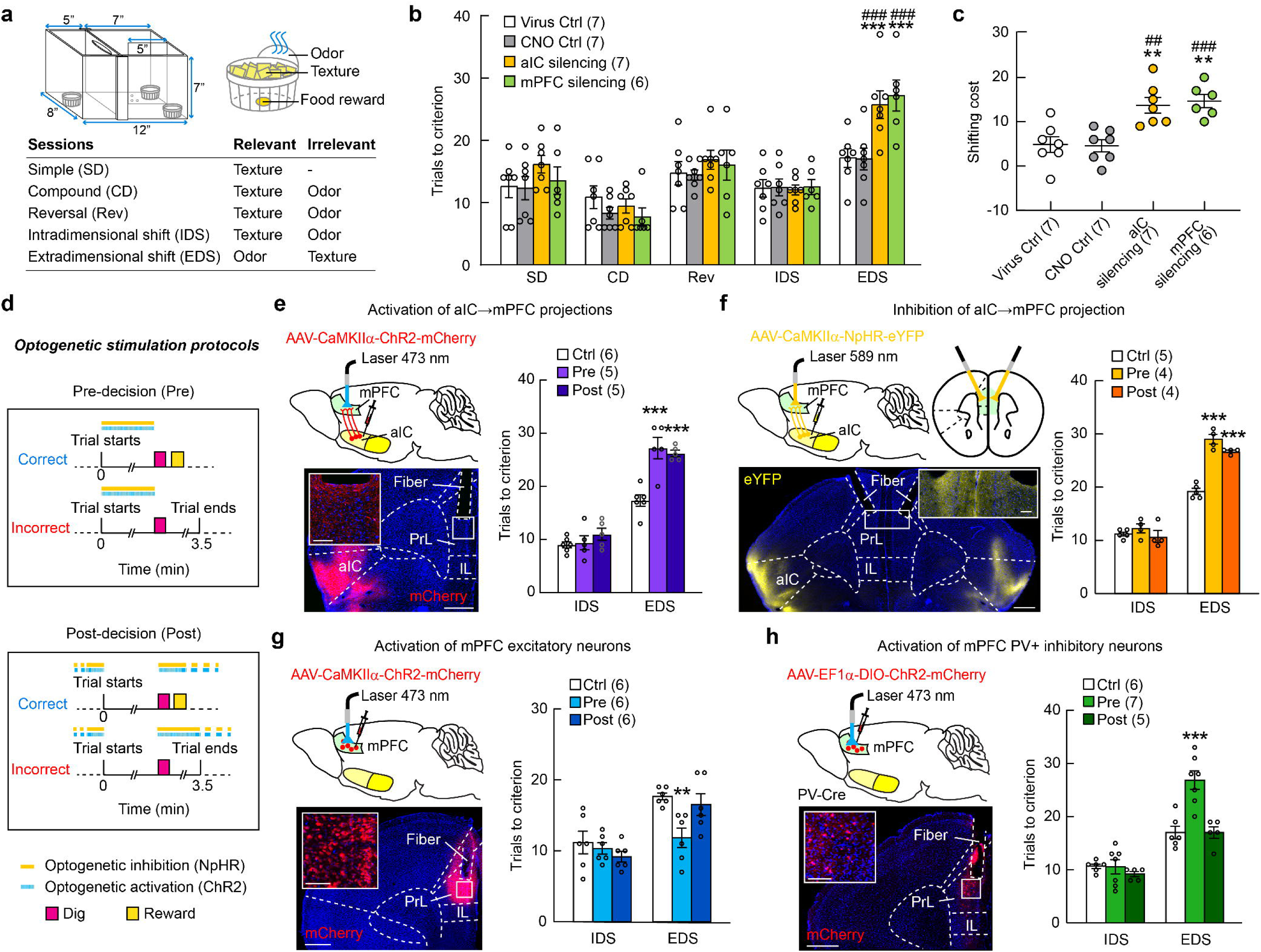
The aIC→mPFC circuit plays a crucial role in cognitive flexibility. **a,** Schematic of the AST setup and the configuration of AST sessions with reward-relevant and reward-irrelevant sensory cues. **b,c,** Pharmacogenetic inactivation of either mPFC or aIC excitatory neurons impairs EDS performance, but does not affect SD, CD, Rev and IDS. **b**, Trials to criterion in each session. Two-way repeated measures ANOVA followed by Dunnett’s multiple comparisons. Main effect for Treatment: *F*(3,23) = 2.543, *p* = 0.0812; main effect for Session: *F*(4,92) = 38.85, *p* < 0.0001; interaction between Treatment and Session: *F*(12,92) = 2.684, *p* = 0.0039. Post hoc testing indicates that in EDS session, both aIC and mPFC silencing result in significantly more trials to criterion than virus Ctrl (*p* = 0.0008 and 0.0001, respectively), as well as than CNO Ctrl (*p* = 0.0007 and 0.0001, respectively). **c**, Shifting cost (the difference in trials to criterion between IDS and EDS). One-way ANOVA followed by Dunnett’s multiple comparisons. Main effect for Treatment: *F*(2,23) = 0.1050, *p* < 0.0001. Post hoc testing indicates that both aIC and mPFC silencing result in significantly higher shifting cost than virus Ctrl (*p* = 0.0019 and 0.001, respectively), as well as than CNO Ctrl (*p* = 0.0014 and 0.0008, respectively). *: compared to Virus Ctrl, ^#^: compared to CNO Ctrl (**b,c**). **d,** Protocols of pre- and post-decision optogenetic stimulation. **e,f**, Both pre- and post-decision optogenetic activation (**e**) and inhibition (**f**) of aIC axonal terminals in mPFC impair EDS performance. Left: schematic of virus injection and optical fiber implantation, and *post hoc* histology showing the optical fiber tract in mPFC and virus injection site in aIC. The inset shows an enlarged view of aIC→mPFC axonal terminals (red or yellow) in the white box. Right: Trials to criterion in IDS and EDS under different experimental conditions. Two-way repeated measures ANOVA followed by Dunnett’s multiple comparisons with Ctrl. **e**, Main effect for Treatment: *F*(2,13) = 14.88, *p* = 0.0004; main effect for Session: *F*(1, 13) = 195.1, *p* < 0.0001; interaction between Treatment and Session: *F*(2,13) = 8.637, *p* = 0.0041, post hoc testing indicates that in EDS session, both Pre-decision and Post-decision optogenetic activation of aIC→mPFC projections result in significantly more trials to criterion than Ctrl (*p* < 0.0001 for both). **f**, Main effect for Treatment: *F*(2,10) = 30.7, *p* < 0.0001; main effect for Session: *F*(1,10) = 554.9, *p* < 0.0001; interaction between Treatment and Session: *F*(2,10) = 25.37, *p* = 0.0001, post hoc testing indicates that in EDS session, both Pre-decision and Post-decision optogenetic inhibition of aIC→mPFC projections result in significantly more trials to criterion than Ctrl (*p* < 0.0001 for both). **g,** Pre-decision optogenetic activation of mPFC excitatory neurons improves EDS performance. Left: Schematic of virus injection and optical fiber implantation, and *post hoc* histology showing virus-labeled mPFC neurons; Right: Trials to criterion in IDS and EDS under different experimental conditions. Two-way repeated measures ANOVA followed by Dunnett’s multiple comparisons with Ctrl. Main effect for Treatment: *F*(2,15) = 4.789, *p* = 0.0246; main effect for Session: *F*(1,15) = 25.16, *p* = 0.0002; interaction between Treatment and Session: *F*(2,15) = 3.196, *p* = 0.0698, post hoc testing indicates that in EDS session, Pre-decision optogenetic activation of mPFC excitatory neurons result in significantly fewer trials to criterion than Ctrl (*p* = 0.0026). **h,** Pre-decision optogenetic activation of mPFC PV+ inhibitory interneurons impairs EDS performance. Left: Schematic of virus injection and optical fiber implantation, and *post hoc* histology showing virus-labeled mPFC neurons. Right: Trials to criterion in IDS and EDS under different experimental conditions. Two-way repeated measures ANOVA followed by Dunnett’s multiple comparisons with Ctrl. Main effect for Treatment: *F*(2,15) = 19.87, *p* < 0.0001; main effect for Session: *F*(1,15) = 73.31, *p* < 0.0001; interaction between Treatment and Session: *F*(2,15) = 7.564, *p* = 0.0054, post hoc testing indicates that in EDS session, Pre-decision optogenetic activation of PV+ neurons in mPFC result in significantly more trials to criterion than Ctrl (*p* < 0.0001). Data are presented as mean ± s.e.m.. Sample sizes (number of mice) are given in the graphs. **,^##^*p*<0.01, ***,^###^*p*<0.001. aIC: anterior insular cortex, PrL: prelimbic cortex, IL: infralimbic cortex; Scale bars (**e-h**): 500 μm, 100 μm (insets).

**Table 1.**
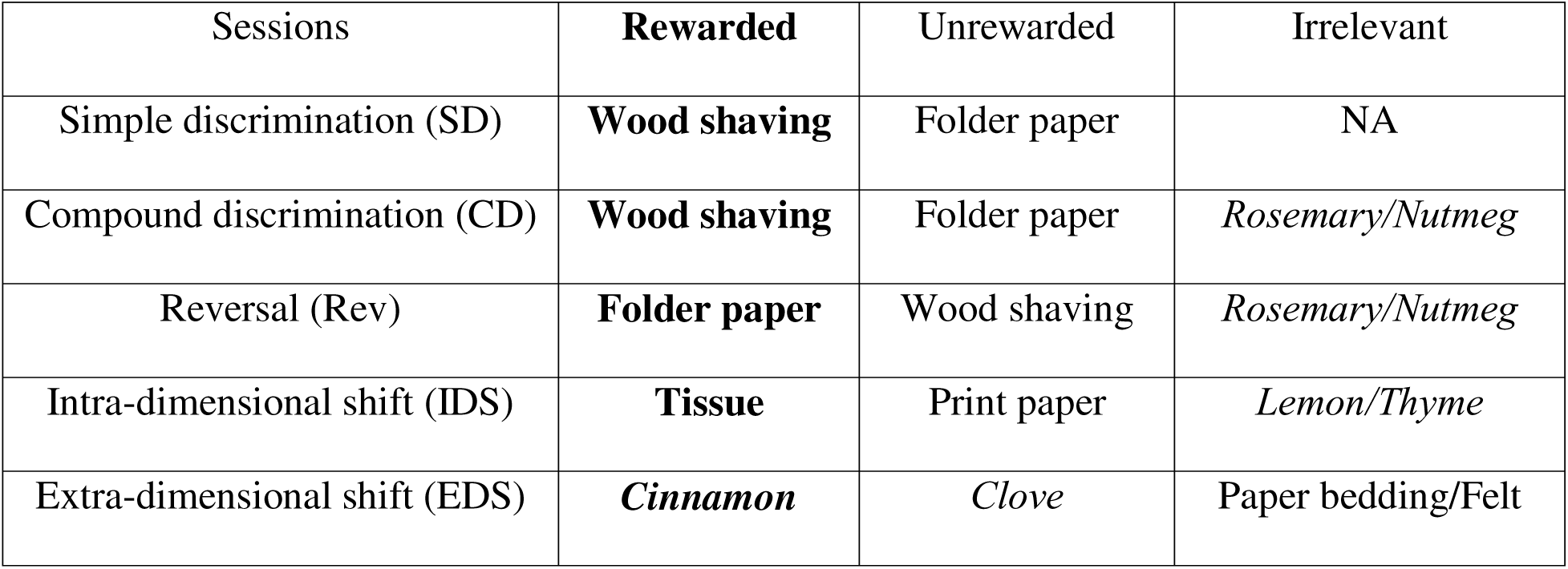
An example of the set of digging media and odors used in different sessions of AST. Rewarded exemplars are in bold. Odors are in italic.

| Sessions | **Rewarded** | Unrewarded | Irrelevant |
| --- | --- | --- | --- |
| Simple discrimination (SD) | **Wood shaving** | Folder paper | NA |
| Compound discrimination (CD) | **Wood shaving** | Folder paper | *Rosemary/Nutmeg* |
| Reversal (Rev) | **Folder paper** | Wood shaving | *Rosemary/Nutmeg* |
| Intra-dimensional shift (IDS) | **Tissue** | Print paper | *Lemon/Thyme* |
| Extra-dimensional shift (EDS) | *Cinnamon* | *Clove* | Paper bedding/Felt |

We found that pharmacogenetic silencing of aIC using a Designer Receptors Exclusively Activated by Designer Drugs (DREADD) system^36^ selectively impairs EDS performance without affecting earlier sessions (Fig. 1b,c and Extended Data Fig. 1). This was evidenced by an increased number of trials to reach criterion in EDS (Fig. 1b) and a higher shifting cost (the difference in trials to criterion between EDS and IDS, Fig. 1c). These impairments resemble the effects following mPFC silencing (Fig. 1b,c). In contrast, performance in earlier AST sessions was unaffected, suggesting that the cognitive demands unique to cross-modality set-shifting in EDS, rather than general sensory discrimination or reward-associated motor responses, critically depend on both mPFC and aIC function.

As aIC directly projects to mPFC (Extended Data Fig. 2), we further investigated the role of the aIC→mPFC pathway in EDS by optogenetics. We infected aIC excitatory neurons with an adeno-associated virus (AAV) encoding either excitatory (ChR2) or inhibitory (NpHR) opsins and optogenetically manipulated their axonal terminals in the mPFC during two distinct time windows of EDS trials: pre-decision (from trial start to digging) and post-decision (from digging to the start of the next trial) (Fig. 1d). Notably, both activation and inhibition of the aIC→mPFC pathway impaired EDS performance (Fig. 1e,f). This effect contrasts with mPFC manipulations, in which only pre-decision manipulation altered EDS performance. In addition, mPFC inhibition (by activating parvalbumin-expressing [PV+] inhibitory interneurons) impaired EDS performance, whereas mPFC activation (by activating excitatory neurons) improved it (Fig. 1g,h). Taken together, our findings reveal distinct roles of mPFC neurons and aIC→mPFC projections in AST, highlighting the unique requirement of tightly regulated aIC→mPFC activity following decision-making to ensure optimal set-shifting.

### aIC→mPFC carries outcome information

To further investigate the role of aIC→mPFC neurons in AST, we used fiber photometry to measure their calcium (Ca^2+^) activity during EDS and compared it with the activity of mPFC excitatory neurons. We labeled aIC→mPFC neurons by injecting a retrograde AAV encoding Cre recombinase into the mPFC and a Cre-dependent GCaMP7f-encoding AAV into the ipsilateral aIC. We implanted an optic fiber above the aIC injection site to record Ca^2+^ signals (Fig. 2a and Extended Data Fig. 3a). To image mPFC excitatory neuron activity, we injected an AAV encoding GCaMP6f under the excitatory neuron-specific promoter CaMKIIα directly into the mPFC and implanted an optic fiber above the injection site (Fig. 2b and Extended Data Fig. 3b). Behavioral events, including trial start (door opening), approach to the ramekin, decision (correct dig vs. incorrect dig), outcome (reward consumption [eat] vs. leave), and trial end (return to the resting area), were manually annotated and aligned with Ca^2+^ traces (Fig. 2c). We observed distinct peri-event activity patterns in aIC→mPFC and mPFC neurons. Both exhibited increased activity at trial start, but aIC→mPFC neuron activity decreased at trial outcome, whereas mPFC activity decreased at decision (Fig. 2d,e).

**Figure 2.**
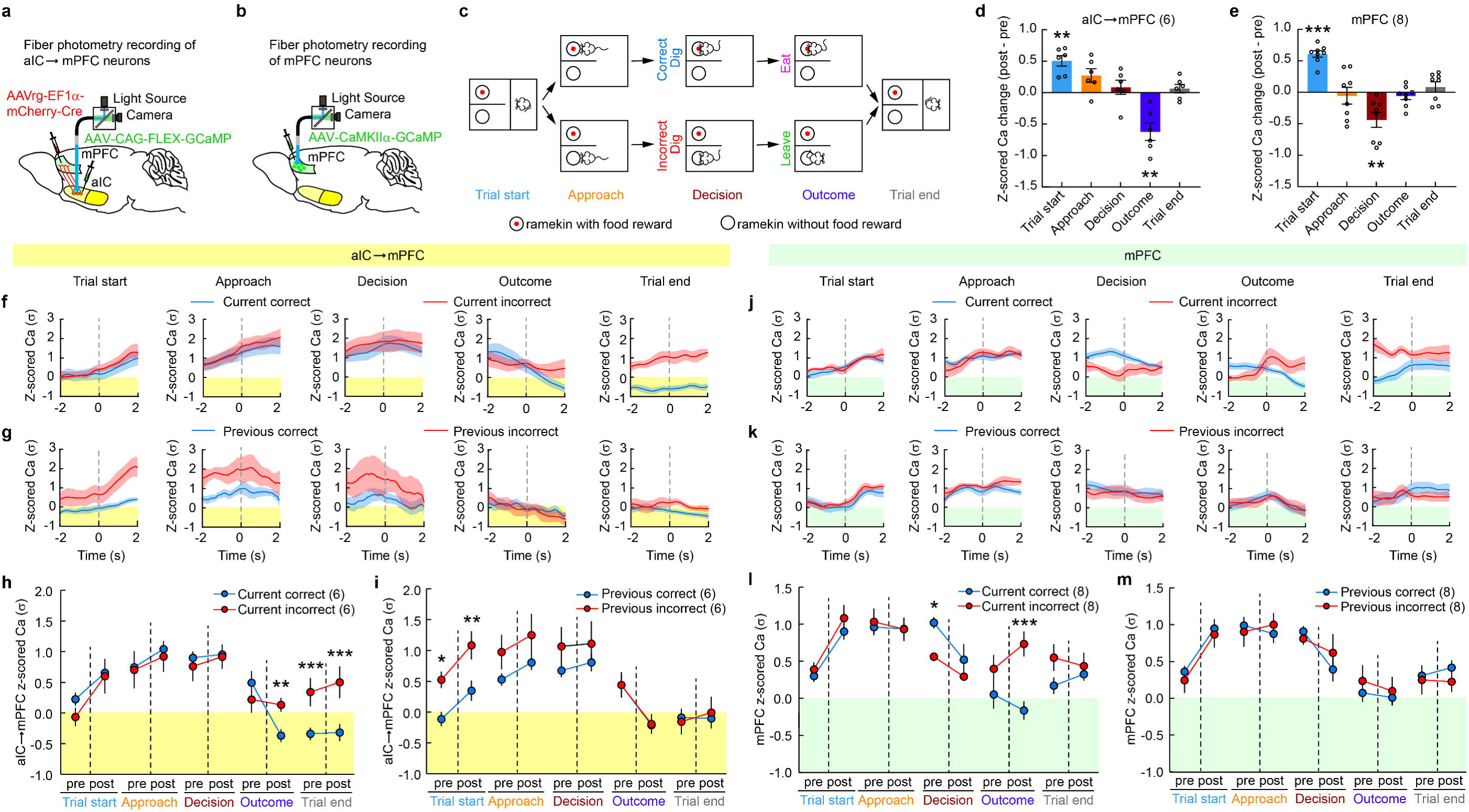
aIC→mPFC neurons exhibit elevated activity following incorrect trials, which persists into subsequent trials. **a,b,** Schematic of virus injection and fiber photometry recording of aIC→mPFC neurons (**a**) and mFPC excitatory neurons (**b**). **c**, Key behavioral events annotated during AST. **d,e,** Ca^2+^ activity changes in aIC→mPFC neurons (**d**) and mPFC neurons (**e**) aligned with key behavioral events. One-sample *t*-test compared to zero. **d**, Trial start: *t*(5) = 6.035, *p* = 0.0018; Approach: *t*(5) = 2.538, *p* = 0.0520; Decision: *t*(5) = 0.7614, *p* = 0.4808; Outcome: *t*(5) = 4.519, *p* = 0.0063; Trial end: *t*(5) = 0.9316, *p* = 0.3943. **e**, Trial start: *t*(7) = 11.99, *p* < 0.0001; Approach: *t*(7) = 0.4015, *p* = 0.7; Decision: *t*(7) = 3.685, *p* = 0.0078; Outcome: *t*(7) = 0.9996, *p* = 0.3508; Trial end: *t*(7) = 1.049, *p* = 0.3289. **f-i**, Ca^2+^ activity in aIC→mPFC neurons separated by current trial’s outcome (**f, h)** or previous trial’s outcome (**g, i**). **f**,**g**, Representative Ca^2+^ traces from one mouse. Solid lines represent the mean values of specific trials, shadows are s.e.m. **h,i**, Average of 6 mice. Two-way repeated measures ANOVA followed by Sidak’s multiple comparisons. **h**, Main effect for Decision type: *F*(1,5) = 0.4684, *p* = 0.5242; main effect for Behavioral event: *F*(9,45) = 15.48, *p* < 0.0001; interaction between Decision type and Behavioral event: *F*(9,45) = 8.517, *p* < 0.0001. Post hoc testing indicates that Post Outcome, Pre Trial end and Post Trial end show significant difference between correct and incorrect trials (*p* = 0.0076, *p* = 0.0001, *p* < 0.0001, respectively). **i**, Main effect for Decision type: *F*(1,5) = 2.462, *p* = 0.1774; main effect for Behavioral event: *F*(9,45) = 17.54, *p* < 0.0001; interaction between Decision type and Behavioral event: *F*(9,45) = 2.073, *p* = 0.0525. Post hoc testing indicates that Pre Trial start and Post Trial start show significant difference between previous correct and previous incorrect trials (*p* = 0.0187, *p* = 0.0043, respectively). **j-m**, Ca^2+^ activity in mPFC neurons separated by current trial’s outcome (**j,l**) or previous trial’s outcome (**k,m**). **j,k** Representative Ca^2+^ traces from one mouse. Solid lines represent the mean values of specific trials, shadows are s.e.m.. **l,m**, Average of 8 mice. Two-way repeated measures ANOVA followed by Sidak’s multiple comparisons. **l**, Main effect for Decision type: *F*(1,7) = 1.646, *p* = 0.2404; main effect for Behavioral event: *F*(9,63) = 11.37, *p* < 0.0001; interaction between Decision type and Behavioral event: *F*(9,63) = 5.513, *p* < 0.0001. Post hoc testing indicates that Pre Decision and Post Outcome show significant difference between current correct and incorrect trials (*p* = 0.0431, *p* < 0.0001, respectively). **m**, Main effect for Decision type: *F*(1,7) = 0.001, *p* = 0.9730; main effect for Behavioral event: *F*(9,63) = 8.062, *p* < 0.0001; interaction between Decision type and Behavioral event: *F*(9, 63) = 1.012, *p* = 0.4403. Post hoc testing reveals no significant difference between previous correct and previously incorrect trial at any behavioral events. Data are presented as mean ± s.e.m.. Sample sizes (number of mice) are given in the graphs. \**p*<0.05, \*\**p*<0.01, \*\*\**p*<0.001.

After separating correct and incorrect trials, we found that aIC→mPFC neurons exhibited higher activity following the outcome of incorrect trials compared to correct ones, with this heightened activity persisting into the next trial (Fig. 2f-i and Extended Data Fig. 3c-f). In contrast, mPFC neurons displayed higher pre-decision activity and lower post-outcome activity during correct trials compared to incorrect trials. This difference dissipated by the end of the trial, and prior outcomes did not influence mPFC activity in subsequent trials (Fig. 2j-m and Extended Data Fig. 3g-j). These findings highlight the unique task engagement of aIC→mPFC neurons, characterized by outcome-dependent activity that persists across trials.

To determine whether the dynamics of mPFC and aIC→mPFC neural activities identified above are unique to EDS, we analyzed neural responses during Rev, which maintains the reward-relevant sensory modality but reverses the cue-reward association. We selected Rev over other sessions because SD includes only one sensory modality, while CD and IDS have substantially fewer trials. We found that both mPFC and aIC→mPFC projection neurons exhibited peri-event dynamics in Rev similar to those observed in EDS (Extended Data Fig. 4). Although mPFC is not required for reversal learning, the conserved dynamics observed in both Rev and EDS sessions suggest that mPFC neurons and aIC→mPFC projections serve general roles in encoding decision- and outcome-related information. The functional necessity of this circuit, however, becomes evident only when cross-modal rule shifts demand flexible updating of decision strategies.

### Perturbing aIC→mPFC disrupts mPFC update

How do aIC inputs affect mPFC neuronal activity patterns? To answer this question, we imaged mPFC neurons during AST, with and without optogenetic activation of aIC→mPFC projections during the pre-decision phase of EDS sessions (Fig. 3a-d). Optogenetic manipulation of aIC→mPFC projections did not affect locomotion, nor did it affect approach latency, approach-to-dig time, or food consumption during EDS (Extended Data Fig. 5). Aligning Ca^2+^ traces with the behavioral video, we first identified neurons modulated by key events (trial start, approach, decision, and outcome) in EDS trials (Extended Data Fig. 6a). The fraction of mPFC neurons selectively modulated by all behavioral events was comparable between control and optogenetic manipulation (Extended Data Fig. 6b), suggesting that activation of aIC→mPFC projection did not broadly alter task-related neuronal activity.

**Figure 3.**
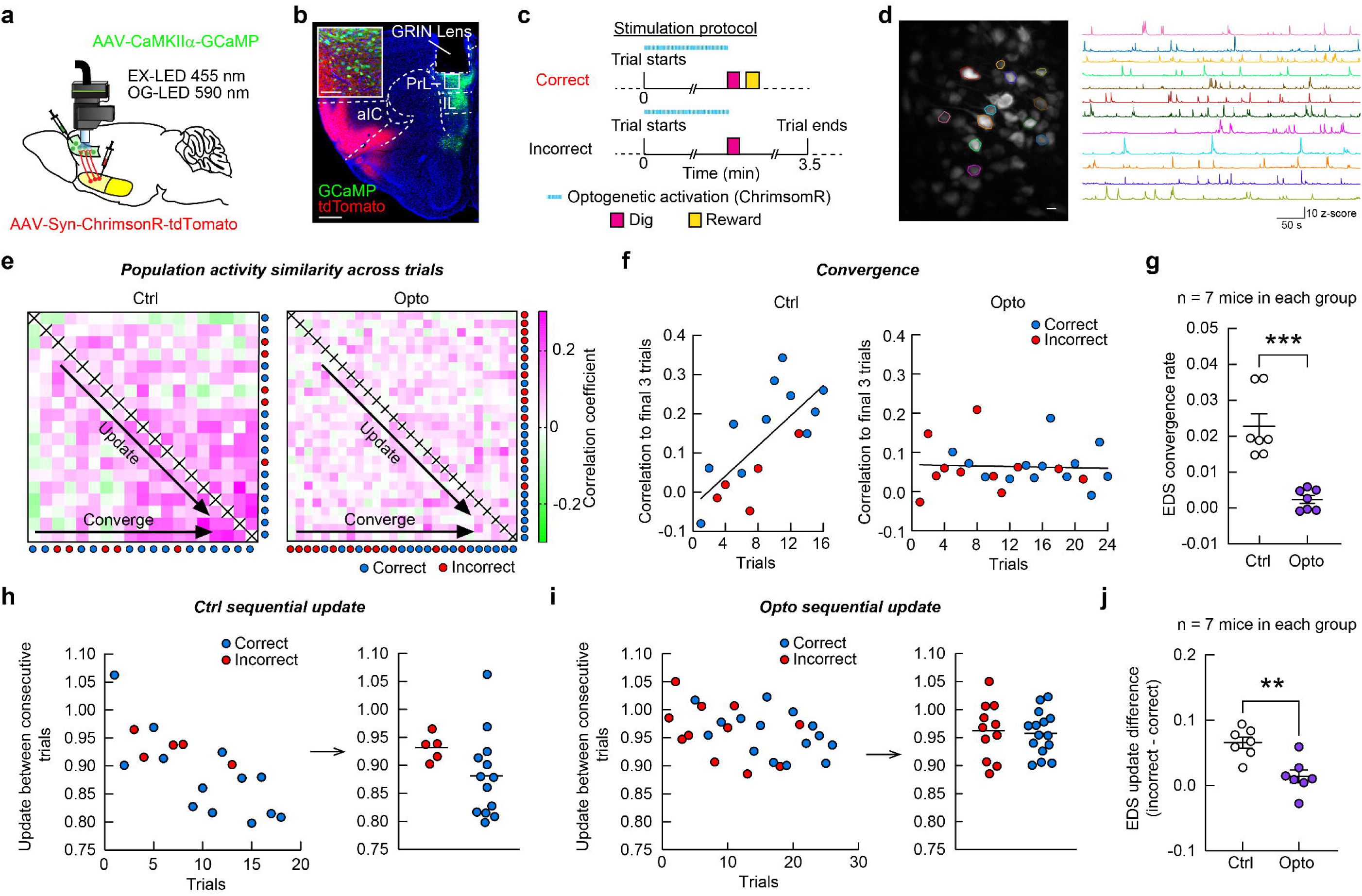
Manipulation of aIC→mPFC projection prevents outcome-specific update and convergence of mPFC population activity patterns during EDS. **a,** Schematic of virus injection and miniscope implantation. EX-LED: excitation LED. OG-LED: optogenetic LED. **b**, *Post hoc* histology showing virus expression in mPFC and aIC, and the GRIN lens tract over mPFC. Inset: Enlarged view of the white box showing GCaMP+ excitatory neurons (green) and aIC synaptic terminals expressing ChrimsonR-tdTomato (red) in the mPFC. Scale bars: 500 μm, 100 μm (inset). **c**, Protocol of pre-decision optogenetic stimulation, performed in conjunction with miniscope imaging. **d**, Representative miniscope Ca^2+^ image after preprocessing and maximum intensity projection along with z-scored Ca^2+^ traces extracted from the neurons labeled in the left image. Scale bar: 20 μm. **e**, Correlation matrices showing pairwise comparisons of mPFC population activity patterns during EDS trials in two example mice. Ctrl: control mouse; Opto: mouse with optogenetic stimulation. Each matrix element represents the Pearson correlation coefficient between a pair of population activity patterns. Pairwise similarity increases in late trials in Ctrl but not Opto mouse. **f**, The correlation between each EDS trial’s population activity pattern and the final pattern (average of the last three trials) increases with trial progression in the example Ctrl mouse (left) but not in the Opto mouse (right). The slope of the regression line relating the correlation coefficients to the trial number indicates the rate of convergence. **g**, Optogenetic stimulation of the aIC→mPFC projection significantly reduces the convergence rate of mPFC population activity patterns during EDS trials compared to the control condition across animals. Unpaired t-test, *t*(12) = 5.544, *p* = 0.0001, n = 7 mice per condition. **h,** Incorrect outcomes induce greater update of mPFC population activity patterns in the next trial compared to correct outcomes in the example Ctrl mouse. The update rate is quantified as the distance (*d*) between the population activity patterns in consecutive trials, *d* = 1 - *ρ*, where ρ is the Pearson correlation coefficient as defined in (**e**). Left: mPFC updates plotted against trial numbers; Right: the average update rates (horizontal lines) following correct vs incorrect trials. **i**, Trial-by-trial updates from the example Opto mouse. Left: mPFC updates plotted against trial numbers; Right: the average update rates (horizontal lines) following correct vs incorrect trials. **j**, Pre-decision optogenetic stimulation of the aIC→mPFC projection decreases the trial-outcome dependence of mPFC updating. Unpaired t-test, *t*(12) = 3.993, *p* = 0.0018, n = 7 mice per condition. Data are shown as mean ± s.e.m. Sample sizes (number of mice) are given in the graphs. \*\**p*<0.01, \*\*\**p*<0.001.

Previous studies have shown that neuronal ensembles in task-relevant cortical regions evolve and converge as animals adapt to new tasks, for example, in the motor cortex during motor learning^37,38^ and in the orbitofrontal cortex (OFC) during odor sequence tasks^39^. Motivated by these studies, we compared mPFC neuronal ensembles during the pre-decision phase (from trial start to digging) across EDS trials. We quantified the similarity of neuronal activity patterns between pairs of trials by calculating the Pearson correlation coefficient. In control mice, this similarity increased progressively over successive trials, with activity patterns from later trials showing greater resemblance to one another (Fig. 3e, left). Importantly, this phenomenon emerges at the ensemble level. While some individual neurons show increasingly consistent activity patterns toward the end of the EDS trials, others remain variable (Extended Data Fig. 7). In contrast, this trend was absent in mice with optogenetic activation of the aIC→mPFC projection during the same period (Fig. 3e, right). To further quantify this effect, we compared activity patterns from earlier trials with the average pattern in the last three trials and found a gradual increase in correlation across trials in control mice, indicating the convergence of neural activity patterns. Pre-decision optogenetic activation of the aIC→mPFC projection significantly disrupted this process, reducing the rate of activity pattern convergence (Fig. 3f,g).

To determine whether the increased correlation in neuronal population dynamics may simply reflect sensory-motor associations, we restricted our analysis to correct EDS trials, which involved the same sensory cues, motor responses, and outcomes. Even among correct trials, mPFC activity patterns in early trials differed significantly from those in later trials (Fig. 3f, Extended Data Fig. 8a,b), suggesting that mPFC population dynamics undergo progressive adaptation rather than reflecting a fixed sensorimotor representation. Increased correlation between trials could, in principle, arise from reduced overall neural activity rather than from structured encoding in neuronal populations. To test this, we identified trial pairs whose correlation values were among the top 5% of all trial pairs (“high-correlation group”), and those among the bottom 5% (“low-correlation group”). We then quantified the mean activity of the neuronal population across all pairs in each group. We found no significant difference in the mean activity between high- and low-correlation groups (Extended Data 8c), suggesting that elevated correlation is more likely driven by structured population co-activation rather than widespread neuronal inactivity. To further validate our findings, we also applied the non-parametric Spearman rank correlation to assess trial-to-trial similarity in neural activity patterns. This analysis yielded conclusions consistent with those obtained using Pearson correlation, supporting the observed pattern of neural convergence (Extended Data Fig. 9).

To better understand how the aIC→mPFC projection influences convergence of mPFC activity, we analyzed trial-to-trial updates in mPFC population dynamics. The update rate, defined as the correlation distance (one minus the correlation coefficient) between consecutive trials, was calculated separately for correct and incorrect trials. In control mice, incorrect trials were followed by larger updates in mPFC activity compared to correct trials (Fig. 3h,i, representative trajectories of population activity shown in Extended Data Fig. 8d. Magenta: first trial in the pair is correct; green: first trial in the pair is incorrect). Optogenetic activation of the aIC→mPFC projection abolished this difference, resulting in similar update rates after correct and incorrect trials (Fig. 3j). Interestingly, neither convergence of mPFC population activity nor outcome-dependent updating was observed in the Rev session of control mice (Extended Data Fig. 10), suggesting that these neural signatures are specific to cross-modality set-shifting and not merely a consequence of general learning or sensorimotor updating.

Given the slow temporal dynamics of calcium signals, we performed an additional analysis by averaging the activity of each cell within four 2-second time windows prior to decision-making (post-trial start, pre-approach, post-approach, and pre-digging). We then repeated the above trial-to-trial correlation analysis using these window-averaged signals. The results were consistent with our original analysis without window-averaging (Extended Data Fig. 11), suggesting that the observed effects are driven by population-level activity patterns rather than precise millisecond-scale event timing. This is also in line with the nature of self-initiated decision-making in AST, where similar neuronal populations are recruited across trials, but the exact timing of their activation relative to behavioral events (e.g., approach, digging) may vary. Together, our findings suggest that the aIC→mPFC projection is critical for outcome-dependent updating and convergence of mPFC activity that underpins cognitive flexibility during set-shifting.

### Targeting aIC→mPFC rescues stress

Restraint stress is a widely used paradigm in animal studies for investigating the effects of stress on the brain and behavior^40^. We subjected adult mice to repeated restraint stress (2 hours per day for 7 consecutive days) and assessed their performance in AST. Stressed mice performed comparably to controls in SD, CD, Rev, and IDS sessions (Fig. 4a), and showed normal approach latency, approach to dig time and food consumption time (Extended Data Fig. 12), indicating that reward learning, motor execution, and reward palatability were intact. However, stressed mice exhibited a selective deficit in EDS, revealing an impairment in shifting attentional set across sensory modalities (Fig. 4a). Fiber photometry recordings further showed that stress abolished the outcome-dependent difference in aIC→mPFC neuronal activity (Fig. 4b,c and Extended Data Fig. 13). Specifically, after incorrect trials, aIC→mPFC activity was significantly reduced in stressed mice compared to controls, whereas no significant difference was observed following correct trials (Fig. 4d).

**Figure 4.**
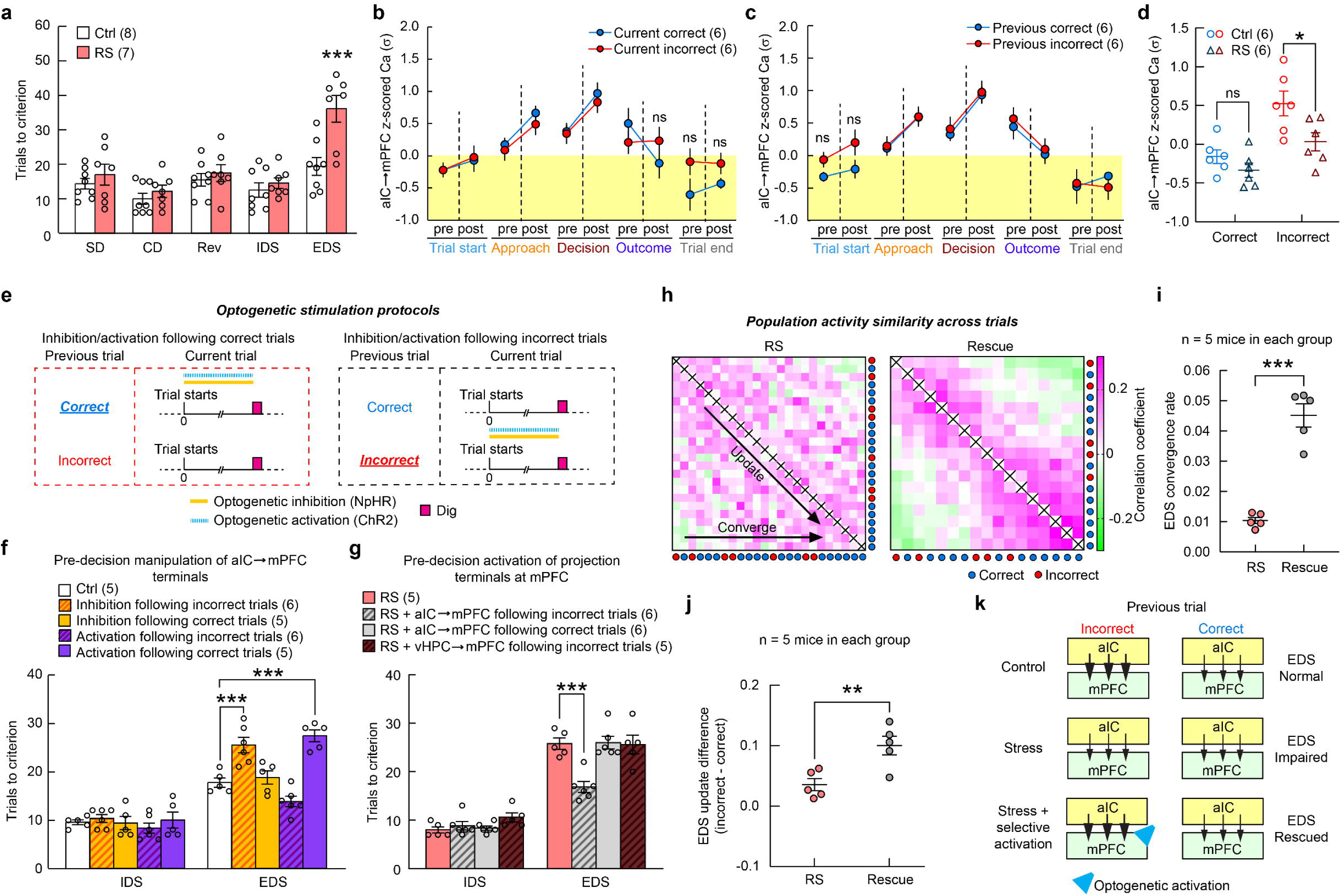
Stress diminishes the outcome-dependence of aIC→mPFC activity, and optogenetic activation following incorrect trials restores cognitive flexibility. **a,** 7-day (7d) restraint stress (RS) selectively impairs EDS. Two-way repeated measures ANOVA followed by Sidak’s multiple comparisons. Main effect for Treatment: *F*(1,13) = 15.18, *p* = 0.0018; main effect for Session: *F*(4,52) = 14.74, *p* < 0.0001; interaction between Treatment and Session: *F*(4,52) = 3.804, *p* = 0.0087. Post hoc testing indicates that 7d RS mice have significantly more trials to criterion than Ctrl mice in EDS session (*p* < 0.0001). **b,c**, Fiber photometry of Ca^2+^ activity in aIC→mPFC neurons separated by current (**b**) or previous (**c**) trial outcomes in 7d RS mice. Two-way repeated measures ANOVA followed by Sidak’s multiple comparisons. **b**, Main effect for Decision type: *F*(1,5) = 0.1476, *p* = 0.7166; main effect for Behavioral event: *F*(9,45) = 14.23, *p* < 0.0001; interaction between Decision type and Behavioral event: *F*(9,45) = 1.743, *p* = 0.1069. Post hoc testing revealed no significant difference between correct and incorrect trials at any behavioral events. **c**, Main effect for Decision type: *F*(1,5) = 2.220, *p* = 0.1964; main effect for Behavioral event: *F*(9,45) = 14.14, *p* < 0.0001; interaction between Decision type and Behavioral event: *F*(9,45) = 0.9622, *p* = 0.4832. Post hoc testing revealed no significant difference between previous correct and previous incorrect trials at any behavioral events. Statistical significance (or lack thereof) between correct and incorrect trials is only indicated for time points that exhibited differences under control conditions. ns: not significant. **d**, Stress decreases the aIC→mPFC neuron activity following incorrect trials. Each data point represents the average neuronal activity of one animal across the five time-windows from post outcome in (**b**) to post trial start in (**c**). Two-way repeated measures ANOVA followed by Sidak’s multiple comparisons. Main effect for Treatment: *F*(1,10) = 4.818, *p* = 0.0529; main effect for Decision type: *F*(1,10) = 62.32, *p* < 0.0001; interaction between Treatment and Decision type: *F*(1,10) = 5.630, *p* = 0.0391. Post hoc testing indicates RS mice have significantly lower aIC→mPFC neurons activity during incorrect trials compared to that of Ctrl mice (*p* = 0.0153). **e**, Protocols of pre-decision optogenetic manipulation of aIC→mPFC axonal terminals following either correct (left) or incorrect trials (right). **f**, Trials to criterion of unstressed mice with or without optogenetic manipulations in IDS and EDS (Optogenetic stimulation only given in EDS). Two-way repeated measures ANOVA followed by Dunnett’s multiple comparisons with Ctrl. Main effect for Treatment: *F*(4,22) = 12.17, *p* < 0.0001; main effect for Session: *F*(1,22) = 247.2, *p* < 0.0001; interaction between Treatment and Session: *F*(4,22) = 10.09, *p* < 0.0001. Post hoc testing indicates that both optogenetic inhibition following incorrect trials and optogenetic activation following correct trials in EDS session led to significantly more trials to criterion than Ctrl (*p* = 0.0002 and *p* < 0.0001, respectively). **g**. Trials to criterion of RS mice with and without projection-specific optogenetic manipulations following incorrect or correct trials in IDS and EDS (Optogenetic stimulation only given in EDS). Two-way repeated measures ANOVA followed by Dunnett’s multiple comparisons with RS. Main effect for Treatment: *F*(3,18) = 10.73, *p* = 0.0003; main effect for Session: *F*(1,18) = 301.7, *p* < 0.0001; interaction between Treatment and Session: *F*(3,18) = 7.967, *p* = 0.0014. Post hoc testing indicates that optogenetic activation aIC→mPFC projections following incorrect trials in EDS session significantly decreases trials to criterion in RS mice (*p* < 0.0001). vHPC: ventral hippocampus. **h**, Representative pairwise comparisons of mPFC population activity patterns during EDS trials in RS and Rescue mouse. Rescue: mouse underwent 7d RS, with pre-decision optogenetic activation of aIC→mPFC axonal terminals after incorrect trials. Each matrix element represents the Pearson correlation coefficient between a pair of population activity patterns. Pairwise similarity increases in late trials in Rescue but not RS mouse. **i**, Significant increase in the convergence rate of mPFC population activity patterns during EDS trials in Rescue mice compared with RS mice. Unpaired t-test, *t*(8) = 8.711, *p* < 0.0001, n = 5 mice per condition. **j**, Rescue mice show significantly higher outcome-specific update of mPFC activity compared to RS mice. Unpaired t-test, *t*(8) = 3.524, *p* = 0.0078, n=5 mice per condition. **k**, Schematic of aIC→mPFC input strength following correct vs incorrect trials and EDS performance under different experimental conditions. Arrow size represents input strength. Data are shown as mean ± s.e.m. Sample sizes (number of mice) are given in the graphs. \**p*<0.05, \*\**p*<0.01, \*\*\**p*<0.001, ns: not significant.

To directly test whether disruption in outcome-dependent signaling of aIC→mPFC projection underlies stress-induced EDS deficits, we optogenetically manipulated aIC terminals in the mPFC in a trial outcome-dependent manner (Fig. 4e). In unstressed mice, inactivation of the aIC→mPFC projection following incorrect trials impaired EDS performance, mimicking the effects of stress, whereas inactivation after correct trials had no effect. Conversely, activation following correct, but not incorrect, trials also disrupted EDS performance (Fig. 4f). These findings suggest that the outcome-specific activity difference in the aIC→mPFC pathway is essential for cognitive flexibility.

We next asked whether restoring this outcome-dependent signal in stressed mice could rescue behavior. Remarkably, optogenetic activation of aIC→mPFC projections specifically after incorrect, but not correct, trials significantly improved EDS performance, restoring it to control levels (Fig. 4g). In contrast, activating ventral hippocampal terminals, another major input to mPFC^41^, using the same protocol failed to improve performance (Fig. 4g), suggesting that the rescue effect is pathway-specific and not due to a general enhancement in mPFC input.

Finally, miniscope imaging of mPFC neurons in stressed mice revealed that trial outcome-specific activation of the aIC→mPFC projection reinstated both the convergence of population activity patterns and the update difference between correct and incorrect trials (Fig. 4h–j). Specifically, selective activation following incorrect trial restored the increase in trial-trial similarity and re-established distinct neural responses following correct versus incorrect outcomes. These ensemble-level changes mirrored the behavioral rescue, supporting a direct link between aIC-driven error signaling and flexible decision-making. Together, these results demonstrate that reinforcing error-related signaling through the aIC→mPFC pathway is sufficient to reverse stress-induced impairment in cross-modal cognitive flexibility (Fig. 4k).

## Discussion

Flexible decision-making depends on the ability to process incorrect outcomes, enabling individuals to learn from mistakes and adapt their actions accordingly^34,42,43^. Errors, inherently surprising events, provide critical salience signals that promote cognitive flexibility. Human imaging studies show that the aIC is activated during perceived errors, suggesting its role in error monitoring and adaptive behavior^44,45^. However, the causal influence of aIC in cognitive flexibility and the underlying neural mechanisms remain unclear. In AST, salience arises when the mouse makes an incorrect choice and thus does not receive an expected reward. We found that aIC→mPFC neurons exhibited a robust, persistent increase in activity following incorrect trials. Notably, this activity emerged only after the outcome was revealed, not during the decision or digging phase, suggesting that it reflects post-decision evaluation rather than the decision process itself. These findings identify a mechanism through which aIC drives adaptive decision-making: by conveying error-related salience signals to mPFC, it updates neural population activity and promotes convergence toward new task rules. This mechanism provides a cellular level account of how aIC translates salience detection into cognitive control, advancing current models of the salience network^2,19^.

While human fMRI studies suggest that the aIC facilitates switching between DMN and CEN^2,18,19^, our findings demonstrate that error-driven aIC→mPFC activity modulates mPFC population dynamics directly. Despite differences in scale and technique, this provides a potential cellular substrate for salience-driven network reconfiguration seen in human studies. Bridging microcircuit mechanisms with large-scale imaging remains a challenge, but our results offer a mechanistic entry point. Future studies combining whole-brain imaging and pathway-specific perturbations will be critical to fully link circuit-level computations to brain-wide adaptive network shifts.

While stress is known to impair cognitive flexibility^13^, its effect on error-related neural signals remains unresolved. Some human electroencephalography (EEG) studies suggest that stress heightens neural responses to mistakes^46^, while others suggest that it dampens them^47^. Both scenarios could hinder learning from mistakes, leading to maladaptive outcomes. However, the limited spatial resolution of EEG and variability in electrode placements prevent direct identification of the underlying neural circuit mechanisms. We showed that stress disrupts the aIC-to-mPFC pathway and dampens salience detection. Specifically, stress abolishes error-related aIC activation, impairing the pathway’s ability to track and integrate incorrect outcomes across trials. This disruption also renders the mPFC update rates after correct and incorrect trials indistinguishable and prevents the convergence of neural activity patterns essential for successful set-shifting. These findings highlight a novel link between stress-induced circuit dysfunction and cognitive rigidity, providing mechanistic insights into stress-related mental disorders where decision-making and adaptability are compromised.

Due to the continuous nature of AST trials, both pre- and post-decision stimulations partially overlap with the post-outcome period from the previous trial. Our photometry data show that aIC→mPFC activity following incorrect outcomes persists into the next trial, suggesting a role in maintaining salience information across trials. Thus, outcome-independent manipulations—whether activation or inhibition—disrupt this ongoing error signal, leading to impaired behavioral flexibility. Excessive activation likely floods the system with an unregulated error signal, while indiscriminate inhibition blocks essential error information, both of which compromise the trial-to-trial updating required for adaptive behavior.

Interestingly, the post-outcome activity of aIC→mPFC neurons persists into the next trial, while mPFC activity is transiently different between correct and incorrect trials. This distinction underscores the unique role of the aIC in maintaining outcome-related information across trials, consistent with its function in error tracking. The persistent activity of the aIC between trials likely supports the integration and transfer of trial outcome signals to the mPFC for adaptive control, whereas the mPFC operates in a more task-phase-specific manner. How does the aIC sustain its activity across the inter-trial interval, which we estimate to average approximately 15 seconds? Previous studies suggest several potential mechanisms for maintaining persistent cortical activity over similar timescales, including intrinsic biophysical properties of neurons, short-term synaptic plasticity, network attractor states, and neuromodulatory inputs^48,49^. Future studies will be needed to clarify how this persistence is maintained and how it interfaces with downstream targets.

Finally, treating impairments in cognitive flexibility in individuals under chronic stress remains challenging, despite extensive efforts through cognitive behavioral therapy and antidepressant-based pharmacotherapy^1,50^. Using an animal model of stress-induced set-shifting deficits, our study demonstrates that impairment of outcome-dependent neural activity in the aIC→mPFC projection plays a critical role in these deficits. Importantly, enhancing activity in aIC→mPFC pathway specifically after incorrect trials is sufficient to restore normal cognitive flexibility. These findings underscore the therapeutic potential of outcome-dependent and pathway-specific neural modulation strategies in overcoming stress-induced rigidity. Given the cross-species applicability of AST and the conserved roles of aIC and mPFC in error monitoring and adaptive control, our findings provide a promising foundation for developing targeted brain stimulation approaches to mitigate cognitive inflexibility in stress-related neuropsychiatric conditions^51^.

## Methods

### Experimental animal

All experiments were conducted in accordance with the National Institutes of Health *Guide for Care and Use of Laboratory Animals* and approved by the Institutional Animal Care and Use Committee of University of California Santa Cruz. C57BL/6J (JAX #000664) and PV-Cre (JAX #008069) mouse line were purchased from The Jackson Laboratory. All mice were maintained on the C57BL/6J background. Mice were group-housed under a 12-hour light-dark cycle with *ad libitum* chow food and water. For all experiment, mice of both sexes (total 103 males and 94 females) between 7-12 weeks of age were used and randomly assigned to different experimental groups.

### Attentional set-shifting task (AST) and behavioral annotation

AST was performed as previously described^52^ with slight modifications. The mouse was handled daily by the experimenter and food restricted to 80-85% of its baseline weight. It was then acclimated to the AST arena and trained to dig for food rewards, which are ∼ 10 mg pieces of Honey Nut Cheerio (General Mills, Minneapolis, MN), in ceramic ramekins (diameter = 2.5”, depth = 1.5”) for 2 days. On the training days, food rewards were placed under increasing amounts of regular bedding materials. On the testing day, the mouse underwent five sessions sequentially: 1) simple discrimination (SD), in which the mouse chose between digging media with two distinct textures, only one being associated with the food reward; 2) compound discrimination (CD), in which two odors were introduced, but the reward was still associated with the same texture as in SD; 3) reversal (Rev), which preserved the same texture/odor combination but the reward-associated texture was swapped; 4) intra-dimensional shift (IDS), which introduced new texture/odor combinations and the reward was still associated with one of the textures; 5) extra-dimensional shift (EDS), which again introduced new texture/odor combinations but the reward was associated with one of the odors rather than textures. An example of the set of textures and odors used in each session is given in Table 1. The positions of the two ramekins are randomized in all trials.

In all sessions, each trial started with the experimenter lifting the gate to allow the mouse to explore the arena and to make a digging choice. As soon as the mouse started digging in a ramekin, the other ramekin was removed from the arena. If the mouse dug correctly, it was allowed to finish consuming the food reward before being returned to the waiting area. If the mouse dug incorrectly, it was left to explore the arena until 3 min from trial start. If the mouse did not dig within 3 min from trial start (which was rare), the trial was considered “no choice”. Each session ended when the mouse completed 6 consecutive trials successfully. The number of trials to criterion and shifting cost (the difference of trials to criterion between IDS and EDS) were used to evaluate the performance. The establishment of an attentional set was demonstrated by the fact that control mice took significantly more trials to finish EDS than IDS.

The behavior of the mouse during AST was recorded using a CMOS camera (acA1300-60gm, Basler AG, Ahrensburg, Germany; 1280 × 960 pixels, frame rate 30 Hz). Key behavioral events in AST were annotated manually using BORIS (v.7.10.2)^53^. “Trial start” was defined as the moment of gate-lifting. “Approach” was defined as the moment when the mouse put its front paw or head over the rim of the ramekin and was exposed to the digging material and odor. The onset of “Correct dig” and “Incorrect dig” was defined as the initiation of digging. The onset of “Eating” was defined as the onset of reward consumption. “Leave” was defined as the moment when the mouse left the ramekin after an incorrect dig.

### Restraint stress

Restraint stress (RS) was induced by placing the mouse in a perforated 50 mL conical tube for 2 h daily, as previously described^54^. Littermates were randomly assigned to the stress or the control group.

### Virus injection

The mouse was anesthetized with isoflurane (4% for induction, 1.5% for maintenance) and mounted on a stereotaxic frame (Angle Two, Leica Biosystems, Deer Park, IL). Ophthalmic ointment was applied to prevent eye desiccation and irritation; dexamethasone (2 μg/g bodyweight) was injected intramuscularly, and carprofen (5 μg/g bodyweight) was injected intraperitoneally. The scalp was disinfected with Alcohol Prep Pad (B60307, PDI, Orangeburg, NY). The skull was exposed by a midline incision in the scalp. A small hole was made through the skull over the target region with a high-speed micro-drill (Foredom K1070, Blackstone Industries, LLC, Bethel, CT). A pulled glass micropipette loaded with viruses (see below for details) was stereotactically lowered through the drilled hole to the desired depth. After 5 min waiting period, injection proceeded at 100 nl/min using a custom-made injector based on a single-axis oil hydraulic micromanipulator (MO-10, Narishige, Tokyo, Japan), followed by a 10 min waiting period before the micropipette was slowly retracted. Additional implantation procedures were performed for optogenetics, fiber photometry, or miniscope imaging preparations (see below). Finally, the wound was closed, and the mouse was put on a warm blanket for recovery before returning to the housing cage. Enrofloxacin (5 μg/g bodyweight) and buprenorphine (0.3 μg/g bodyweight) were injected subcutaneously to prevent infection and pain.

### Anterograde and retrograde viral tracing

We performed anterograde viral tracing by stereotactically injecting AAV2/1-CAG-GFP (2.4E13 gc/ml, 0.3 μl) into the aIC (AP +1.90 mm, ML 2.8 mm, DV −-2.35 mm below dura), and retrograde viral tracing by injecting AAVrg-EF1α-mCherry (1.3E13 gc/ml, 0.3 μl) into the mPFC (AP +1.78 mm, ML 0.3 mm, DV −1.35 mm below dura). For ventral hippocampus (vHPC), we used the coordinate AP −3.1 mm, ML 3.0 mm, DV 4.2 mm. The same coordinates were used for fiber photometry, optogenetics, miniscope imaging, and virus-based tracing experiments unless otherwise noted. After 3 weeks of incubation, we perfused the mouse for histological examination (see the section *Post hoc* histology and immunohistochemistry for details).

### Rabies virus-mediated retrograde trans-synaptic tracing

We used rabies virus-based retrograde tracing to identify the aIC neurons that directly synapse onto excitatory neurons or PV+ INs in mPFC. To label the input neurons to mPFC excitatory neurons, we injected a mixture of AAV2/1-CaMKII0.4-Cre-SV40 (1.9E13 gc/ml, 0.1 μl) and AAV2/8-hSyn-FLEX-TVA-P2A-eGFP-2A-oG (3E13 gc/ml, 0.2 μl) unilaterally into the mPFC of wild-type mice to label the excitatory starter cells; after three weeks of incubation, EnvA-RV-ΔG-dsRed (3E8 IFU/mL, 0.1 μl) was injected into the same region. To label the input neurons to mPFC PV+ INs, we injected AAV2/8-hSyn-FLEX-TVA-P2A-eGFP-2A-oG (3E13 gc/ml, 0.2 μl) unilaterally into the mPFC of PV-Cre mice to label the PV+ starter cells; after three weeks of incubation, EnvA-RV-ΔG-dsRed (3E8 IFU/mL, 0.3 μl) was injected into the same region. Mice were sacrificed 7 days after rabies virus injection for histology. Starter cells were identified by their co-expression of eGFP and dsRed; their presynaptic partners were identified by the expression of dsRed only.

### Pharmacogenetic manipulation

To silence aIC or mPFC excitatory neurons, AAV2/8-CaMKIIα-hM4D-mCherry (4.2E12 gc/ml, 0.3 μl) was injected into aIC or mPFC bilaterally. After 3 weeks of incubation, the mouse was subjected to AST, with the DREADD agonist clozapine N-oxide (CNO, Enzo Life Sciences, NY; 3 mg/kg of bodyweight) administered intraperitoneally 30 min prior to SD. Mice receiving DREADD virus injection but treated with saline (Virus Ctrl), and mice receiving control virus (AAV2/2-CaMKIIα-mCherry, 4E12 gc/ml, 0.3 μl) injection and treated with CNO (CNO Ctrl), were used as controls.

### Optogenetic manipulation

Four sets of optogenetic experiments were conducted. To activate mPFC excitatory neurons, AAV2/1-CaMKIIα-hChR2(H134R)-mCherry (1E13 gc/ml, 0.3 μl) was injected unilaterally into the mPFC of wild-type mouse. To activate mPFC PV+ INs, AAV2/1-EF1α-DIO-hChR2(H134R)-mCherry-WPRE-HGHpA (1E13 gc/ml, 0.3 μl) was injected unilaterally into the mPFC of PV-Cre mouse. To activate aIC→mPFC or vHPC→mPFC axonal terminals, AAV2/1-CaMKIIα-hChR2(H134R)-mCherry (1E13 gc/ml, 0.3 μl) was injected bilaterally into aIC or vHPC. A fiber-optic cannula (200 μm core, 0.39 NA fiber; RWD Life Science Co., Shenzhen, China) was implanted 0.2 mm above mPFC and secured with dental cement (S371, S398, S396, Parkell, Edgewood, NY). To inhibit aIC→mPFC axonal terminals, AAV2/1-CaMKIIα-eNpHR-eYFP (1.6E13 gc/ml, 0.3 μl) was injected bilaterally into aIC. Two fiber-optic cannulas were implanted 0.2 mm above mPFC with ±26° rotation bilaterally. Control mice were injected with AAVs encoding only fluorescent proteins (without opsin), driven by the same promoter as used in the AAVs encoding opsins for the corresponding experimental group. These animals were exposed to either pre- or post- light stimulation protocols as the experimental mice.

A 473 nm laser (MBL-III-472, 200 mW) and a 589 nm laser (MGL-F-589-100mW BJ00849) from Changchun New Industries Optoelectronics Technology Co., Changchun, China was used to deliver optogenetic activation or inhibition. The laser output was controlled by Arduino Uno (Rev 3). The laser intensity was set to 1-5 mW as measured by a photometer (PM100USB, Thorlabs Inc., Newton, NJ) at the output optical fiber end of the ceramic ferrule. A 20 Hz light pulse train (5 ms pulse width) was used to activate mPFC excitatory neurons, aIC→mPFC axonal terminals or vHPC→mPFC axonal terminals, a 40 Hz train (5 ms pulse width) for PV+ INs, and continuous stimulation was used to inhibit aIC→mPFC axonal terminals. Stimulation was applied from trial start to dig (Pre-decision) or from dig to next trial start (Post-decision) during EDS trials. The Previous Correct and Previous Incorrect optogenetic stimulations refer to Pre-decision optogenetic stimulations delivered only when the previous trial was correct or incorrect, respectively, regardless of current decision-making type.

### Fiber photometry Ca recording

We used fiber photometry to measure the activity of mPFC or aIC→mPFC neurons. To label mPFC neurons, AAV2/1-CaMKIIα-GCaMP6f-WPRE-SV40 (2.3E13 gc/ml, 0.3 μl) was injected into mPFC. To label aIC→mPFC neurons, AAVrg-EF1α-mCherry-IRES-Cre (1.3E13 gc/ml, 0.3 μl) was injected into mPFC, and AAV2/1-CAG-FLEX-jGCaMP7f-WPRE (1.5E13 gc/ml, 0.3 μl) was injected into ipsilateral aIC. Following viral injection, a fiber optic cannula was implanted 0.2 mm above the injection site in mPFC or aIC and secured with dental cement. Three weeks after the surgery, the mouse was acclimated to the AST arena and trained to dig in the ramekins, with the fiber photometry apparatus connected. The mouse was then tested on AST with simultaneous fiber photometry (version 4.1, ThinkerTech Bioscience, Nanjing, China). To accommodate the head-mounted recording apparatus, bigger ramekins (diameter = 3.5”, depth = 1.8”) and extended trial duration (3.5 min) were used; the same conditions applied to optogenetics and miniscope imaging experiments. All recordings were made at 50 Hz frame rate with 480 nm LED illumination. Concurrently, the behavior of the mouse was videotaped at 30 Hz frame rate with a camera (acA 1300-60gm, Basler, Inc., Exton, PA) mounted above the arena; videotaping and photometry were coordinated by custom-written scripts in Bonsai (v2.6.3)^55^ through Arduino Uno (Rev 3). The background signal was automatically subtracted by the Thinker Tech software. Briefly, in addition to selecting a region of interest (ROI) corresponding to the optic fiber for fluorescence recording, a background ROI positioned outside the fiber ROI in a fluorescence-free region but within the CMOS camera-recorded field was chosen during recording. Subsequent analysis was performed with custom-written scripts in MATLAB R2022a (MathWorks, Natick, MA). The exported Ca^2+^ trace was corrected for baseline decay by fitting with either a linear polynomial curve or a two-term exponential model based on visual inspection of the whole trace. Then the z-score was calculated from the mean and standard deviation throughout the whole recording session. Manually annotated behavioral events were aligned with z-scored Ca^2+^ traces; the average values within the pre-event time window (−2 s to 0 s) and the post-event time window (0 s to +2 s) were calculated for each trial.

### Miniscope Ca imaging with optogenetic manipulation

We conducted miniscope Ca imaging of mPFC neurons with and without optogenetic activation of aIC→mPFC axonal terminals. AAV2/1-CaMKIIα-GCaMP6f-WPRE-SV40 (2.3E13 gc/ml, 0.3 μl, diluted 1:2 in saline) was injected into mPFC unilaterally, and AAV2/9-Syn-ChrimsonR-tdTomato (2.6E13 gc/ml, 0.3 μl) was injected into aIC bilaterally. An integrated GRIN Lens (1 mm diameter, 4 mm length; ProView Integrated Lens, Inscopix, Mountain View, CA) was implanted 0.2 mm above the injection site of mPFC and secured with dental cement. After 3-4 weeks of recovery and virus incubation, *in vivo* imaging in the free-moving mouse was performed with a head-mounted miniscope (nVoke 2.0, Inscopix). All recordings were made at 20 Hz frame rate using 455 nm LED excitation (0.1 to 0.6 mW) and 2-8× gain. Optogenetic stimulation of ChrimsonR was achieved by a 590 nm LED integrated in the miniscope at 20 Hz (5 ms pulse width, 2-5 mW). The light trains were started and stopped manually through Arduino Uno (Rev 3) during EDS. Concurrent behavioral videotaping was performed as described above.

The Ca^2+^ traces of individual neurons were extracted using Inscopix Data Processing Software (v1.7.1). Briefly, the raw Ca videos were cropped, spatially and temporally down-sampled, spatially filtered (cutoff: 0.005 to 0.500 pixel^−1^), and motion-corrected using the mean image as reference. Individual neurons were identified from the processed Ca videos using the extended constrained non-negative matrix factorization (CNMFe) algorithm^56^. The identified neurons were inspected by the experimenter to exclude overlapping neurons, blood vessels, or fluorescent debris moving in and out of focus. The ΔF/noise traces of neurons were extracted and deconvolved as previously described^57^ before further analysis with custom-written MATLAB programs.

To determine if a neuron was responsive to specific behavioral events (trial start, approach, decision and outcome), we used the receiver operating characteristic (ROC) analysis^58^. For each occurrence of a specific category of behavioral event, we calculated the average of the z-scored Ca signal over a 2 s window before and a 2 s window after the event as the decision variable (DV). We applied a varying threshold to the DV and plotted the true positive rate (TPR) against the false positive rate (FPR) at each threshold to yield an ROC curve. The degree of neural activity modulated by each category of behavioral event was measured by the area under the ROC curve (auROC). For each neuron and each behavior category, the observed auROC was compared to a null distribution of 1,000 auROC values generated from constructing ROC curves over randomly time-shuffled Ca signals. A neuron was considered significantly responsive (a = 0.05) if its auROC exceeded the 95^th^ percentile of the random distribution (auROC < 2.5^th^ percentile considered inhibited response and auROC > 97.5^th^ percentile considered activated response). The rest were classified as unresponsive to behavioral events.

To examine how mPFC neuronal activity patterns change during AST, we first organized the Ca^2+^ activity data in each trial into a 2D matrix (neuron x time). For the time dimension, the activity values from four 2 s time windows (post trial start, pre approach, post approach, and pre dig) were concatenated, which corresponded to the pre-decision period when photo-stimulation was applied. To determine the similarity of neural activity patterns between a pair of trials, the 2D matrix (neuron x time) representation of activity patterns in each trial was vectorized and the Pearson correlation coefficient between the vectors of the two trials was calculated. To quantify the convergence of population activity patterns over trials, we defined the final pattern as the average of the last three trials and computed the correlation between the earlier trial patterns and the final pattern. These correlation coefficients were regressed against the trial numbers, and the linear regression slope represents the convergence rate of neural activity patterns. To quantify the trial-to-trial update of population activity patterns, the update rate between consecutive trials was defined as one minus the correlation coefficient. The update rates were then grouped according to the outcome of the first trial and averaged with each group for comparison between correct and incorrect trials.

To further examine the robustness of our findings, we carried out a series of additional analyses. To determine if higher correlation between trials could be associated with reduced overall neural activity, we identified trial pairs whose Pearson correlation coefficients were among the top 5% of all trial pairs (“high-correlation group”), and those among the bottom 5% (“low-correlation group”), using the activities during the pre-decision phase of each trial. We then quantified the mean population activity across all pairs within each group for each mouse, and compared the difference between the high- and low-correlation groups.

To assess if the update and convergence of population activity patterns depend on the linear relationship captured by Pearson correlation coefficient, we calculated the non-parametric Spearman rank correlation, which captures monotonic relationship between activity patterns. To examine the evolution of activity patterns within correct trials of EDS, we calculated the mean pairwise correlation between the first six correct trials and compared it with that between the last six correct trials. To assess if the correlation changes across trials depend on precise temporal alignment at millisecond time scale, we averaged the neural activity of each cell within four defined 2-second time windows prior to decision-making (post-trial start, pre-approach, post-approach, and pre-dig), then conducted trial-to-trial correlation analysis using these window-averaged signals. To visualize the correlation difference between representative trial pairs, we applied the principal component analysis (PCA) to reduce dimensionality and plotted the trajectories of population activity from two consecutive trials with either high or low correlation in the space spanned by the first two PCA axes.

### Locomotion in the open-field arena

Locomotor activities were examined in an open-field arena^59^. Each mouse was placed in the center of the arena, which is a square area surrounded by white acrylic walls (42 cm × 42 cm), and allowed to explore the chamber for 10 min. A camara (acA1300-60gm, Basler AG, Ahrensburg, Germany, 1280 × 960 pixels, frame rate 30 Hz) mounted from above was used for video recording. The video was analyzed with DeepLabCut (v2.3.11), and the position of the mouse in each video frame was defined as the center of its body. The average speed was calculated. Data analysis was performed by custom-written scripts in MATLAB R2022a (MathWorks, Natick, MA).

### *Post hoc* histology and immunohistochemistry

The mouse was transcardially perfused with 0.1M phosphate-buffered saline (PBS) followed by 4% paraformaldehyde (PFA) in PBS. The brain was removed and post-fixed in 4% PFA overnight at 4°C and then transferred to 30% sucrose solution for 2 days. The brain was coronally sectioned into 40-50 μm slices with a vibratome (VT1000S, Leica Biosystems, Wetzlar, Germany). The brain slices were counterstained with 4’-6-Diamidino-2-phenylindole (DAPI, 300 nM; D3571, Invitrogen, Waltham, MA) and mounted with Fluoromount-G (0110-01, SouthernBiotech, Birmingham, AL).

For cFos immunohistochemistry (IHC), brain slices (40 μm) were blocked with 1× PBS containing 5% normal goat serum (NGS), 5% Bovine Serum Albumin (BSA) and 0.3% Triton X-100 for 2 h at room temperature, and then incubated with rabbit anti-cFos antibody (1:1000; no. 226003, Synaptic Systems, Göttingen, Germany) in 1× PBS containing 5% NGS for 3 days at 4°C. The slices were then washed three times in 1× PBS and incubated with Alexa Fluro 488 goat anti-rabbit secondary antibody (1:500; A-11008, Invitrogen) for 2 h at the room temperature in 1× PBS containing 5% NGS and 0.2% Trion X-100. The slices were mounted as above.

Fluorescence images were taken with an AxioImager Z2 wide-field microscope (Zeiss, Dublin, CA) under a 10× (NA=0.45) air objective. The images were contrast-enhanced in Photoshop CC 2018 (Adobe Inc., Mountain View, CA) for presentation.

### Quantification and statistical analyses

All behavioral and Ca imaging data were analyzed with the analyst blinded to the experimental conditions. All statistical analyses were performed with Prism 9 (GraphPad Software, San Diego, CA) or MATLAB R2022a (MathWorks, Natick, MA). The sample size *n* refers to the number of mice. Unless otherwise indicated, if the sample met the assumptions for parametric tests, we used two-way Analysis of Variance (ANOVA) followed by Sidak’s or Dunnett’s multiple comparisons test. Dunnett’s test was used when comparing multiple experimental groups against a common control group, and Sidak’s test was used when making multiple independent pairwise comparisons). All *post hoc* comparisons were made to controls unless otherwise stated. In the AST behavioral analysis, each session (e.g., Rev, EDS, IDS) was treated as a separate family of comparisons. We report the sample sizes, the statistical tests used, and the *p* values in figure legends. Statistical significance is defined as *p* < 0.05.

## Data availability

The data supporting the findings of this study are available from the corresponding authors upon request.

## Code availability

Custom-written data analysis codes are available upon request.

## Acknowledgements

We thank Zhigang He, Euiseok Kim, Ju Lu, and Chenyan Ma for critical comments on this manuscript, Karen Jimenez Reyes and Isaac M. Lopez for help with behavioral annotation, Leo Hu for help with miniscope Ca data preprocessing, and Benjamin Abrams (UCSC Life Sciences Microscopy Center) for technical support. This work was supported by grants from the National Institutes of Mental Health (R01MH127737 to Y.Z. and K.H.W.; R01MH136381 to Y.Z.), National Institute on Aging (R01AG071787 to Y.Z.), National Institute on Aging (U24AG072701 to K.H.W.) and a Max Planck Fellowship to Y.Z.

## Author Contributions

S.M., K.H.W. and Y.Z. designed the study. S.M. performed the experiments; S.M. and K.H.W. wrote the program for data analysis and performed the analysis. S.M., K.H.W. and Y.Z. wrote the manuscript.

