## Extended Data for "Targeting insulo-frontal pathway to reduce stress-evoked cognitive rigidity"

**Extended Data Figures**

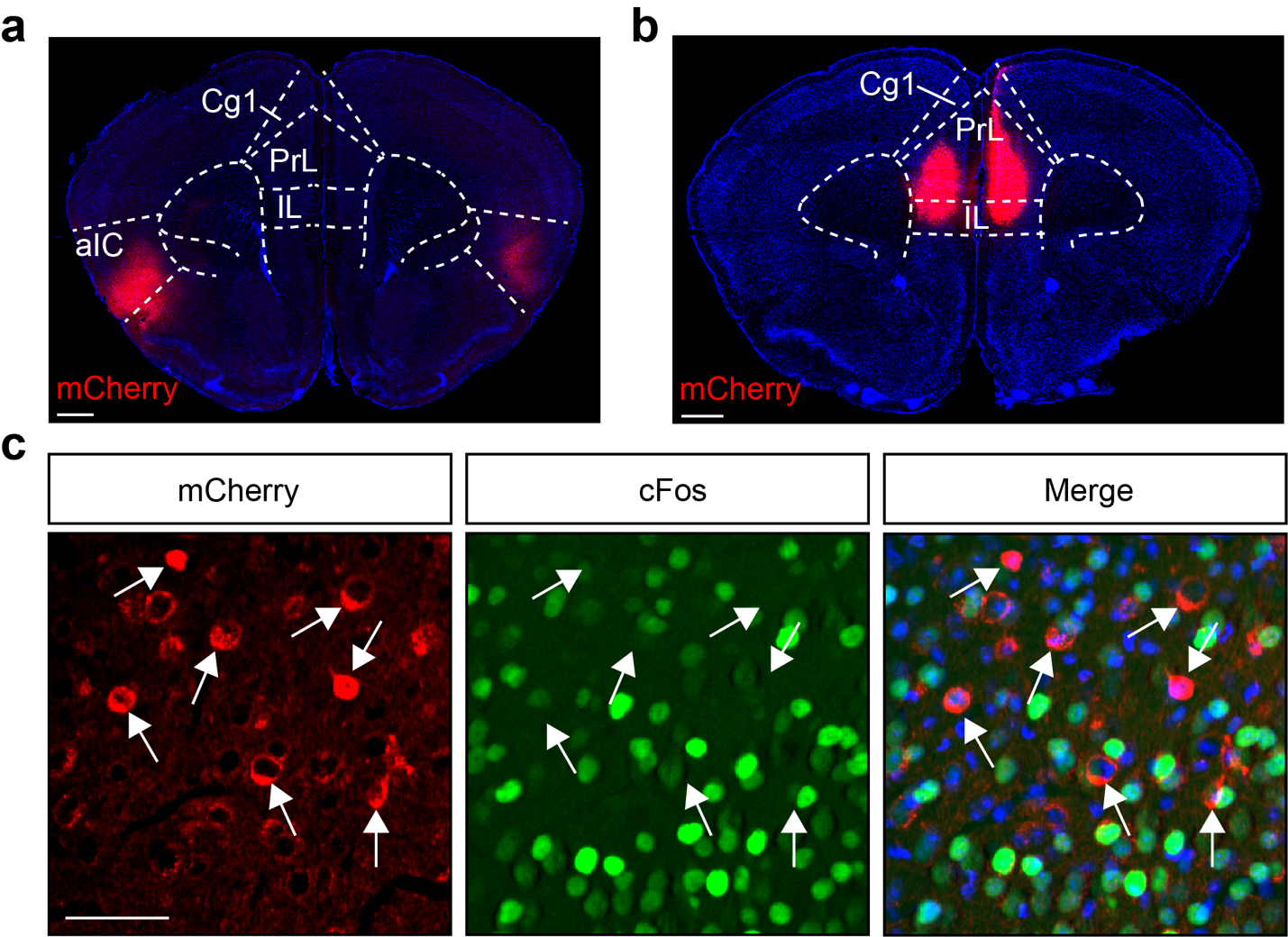

Extended Data Figure 1. Validation of inhibitory DREADD manipulation. a,b, Examples of *post hoc* histological validation of virus injection sites in aIC (a) and mPFC (b). PrL: prelimbic cortex; IL: infralimbic cortex; Cg1: cingulate cortex area 1; aIC: anterior insular cortex. c, Immunohistochemistry shows lower cFos expression level in virus-infected neurons (arrows pointing at representative neurons). Red: mCherry expression by DREADD virus. Green: cFos immunohistochemistry. Blue: DAPI staining. Scale bars: 500 μm (a,b) and 50 μm (c).

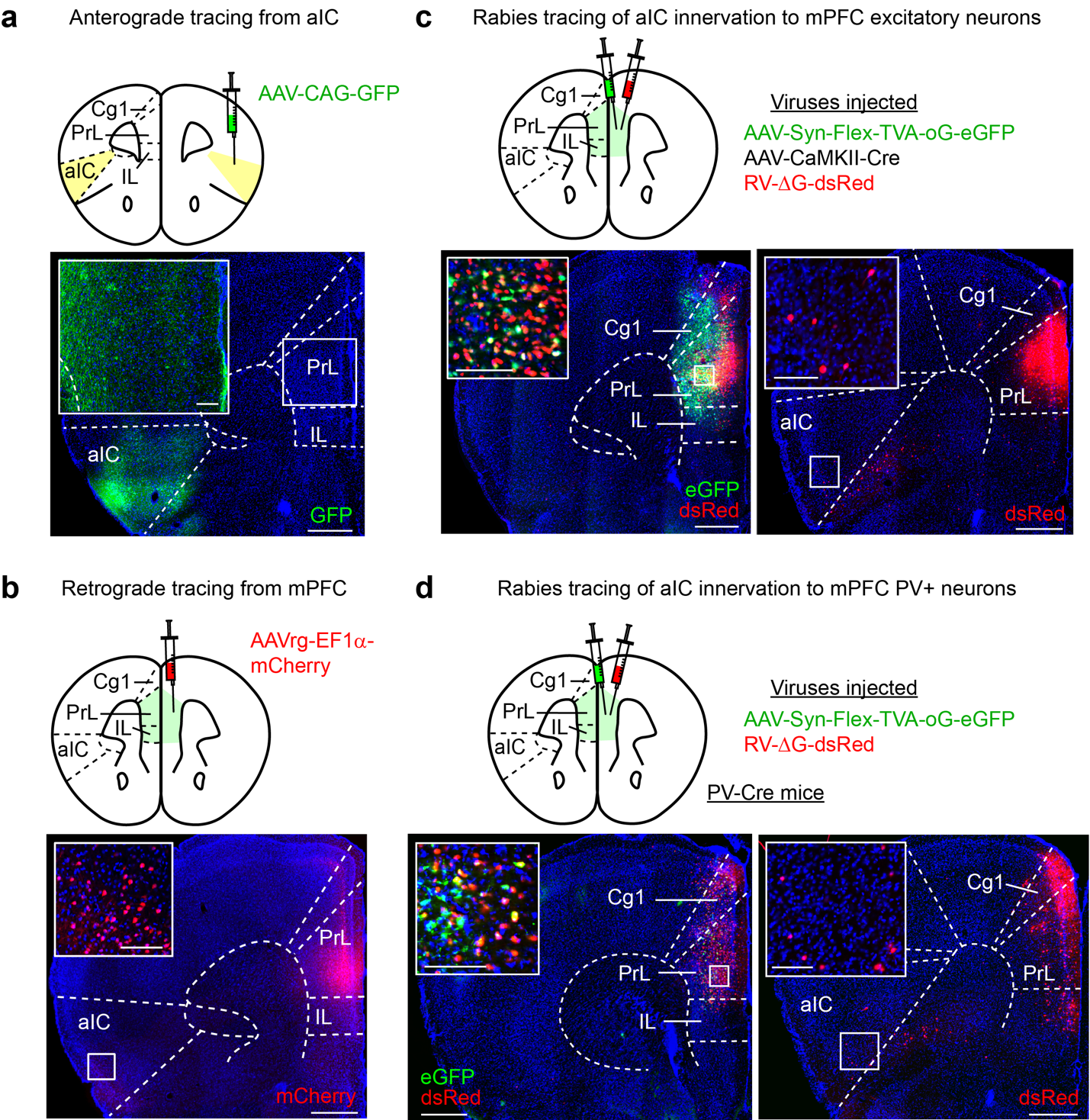

**Extended Data Figure 2. Neurons from aIC innervate both excitatory and inhibitory neurons in mPFC. a,** Anterograde viral tracing reveals aIC neuronal projections in mPFC. Inset: enlarged view of the white box, showing aIC axonal terminals in mPFC. **b**, Retrograde viral tracing from mPFC labels aIC neurons. Inset: enlarged view of the white box, showing aIC neuronal somata. **c,d**, Retrograde trans-synaptic rabies tracing reveals aIC innervation of mPFC excitatory neurons (**c**) and parvalbumin-positive (PV+) inhibitory interneurons (**d**). Insets: enlarged views of the white boxes, showing the injection site (left) and aIC neuronal somata (right). PrL: prelimbic cortex; IL: infralimbic cortex; Cg1: cingulate cortex area 1; aIC: anterior insular cortex. Scale bars: 500 μm, 100 μm (insets).

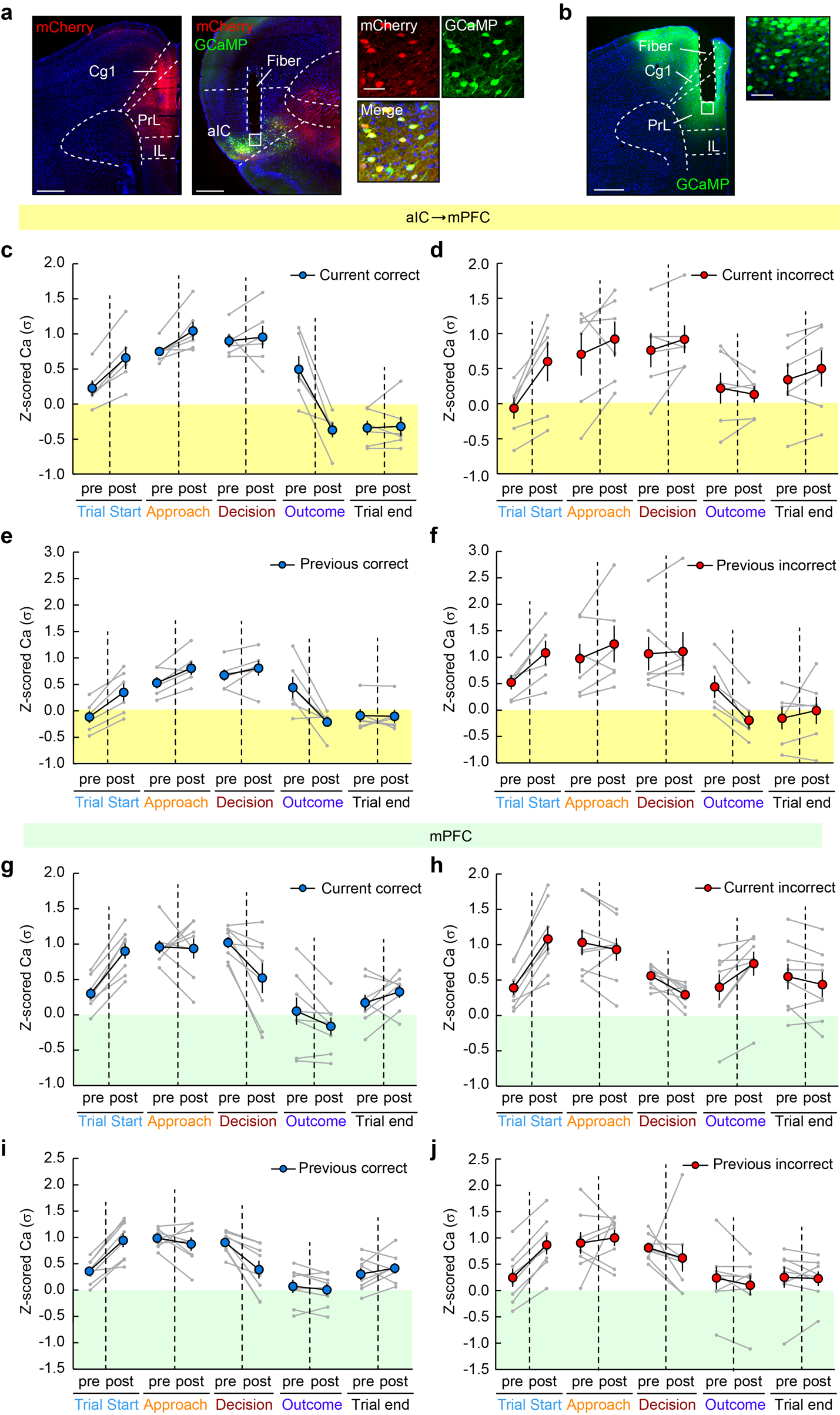

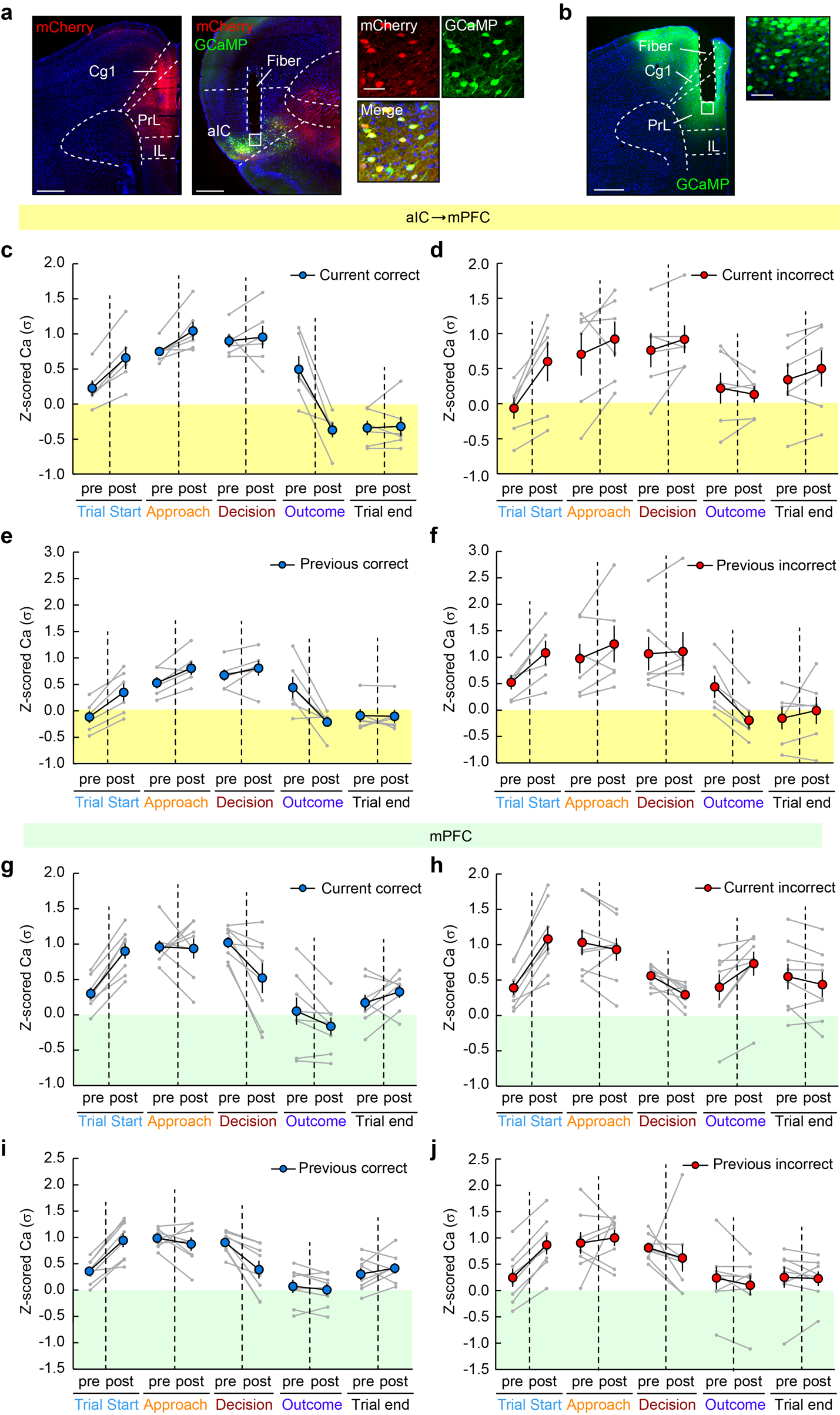

**Extended Data Figure 3. Examples of *post hoc* histological validation of virus injection sites and optical fiber tracts, along with fiber photometry data from individual control mice.** **a**, *Post hoc* histology showing retrograde labeling of aIC→mPFC neurons for fiber photometry recording. Top: AAVrg injection site in mPFC (top left) and projection-specific GCaMP expression, optical fiber tract in aIC (top right). Bottom: enlarged view of the white box, showing retrogradely labeled neurons (mCherry+), GCaMP+ neurons, and their co-localization in aIC. Scale bars: 500 μm (top); 50 μm (bottom). **b**, *Post hoc* histology showing viral labeling of mPFC neurons for fiber photometry recording. Left: optical fiber tract in mPFC; Right: an enlarged view of the GCaMP+ neurons therein. PrL: prelimbic cortex; IL: infralimbic cortex; Cg1: cingulate cortex area 1; aIC: anterior insular cortex. Scale bars: 500 μm (left); 50 μm (right). **c-f**, Ca^2+^ activity of aIC→mPFC neurons in individual animals, separated by current trial’s outcome (**c,d**) or previous trial’s outcome (**e,f**). **g-j**, Ca^2+^ activity of mPFC neurons in individual animals, separated by current trial’s outcome (**g,h**) or previous trial’s outcome (**i,j**). Data are shown as mean ± s.e.m.

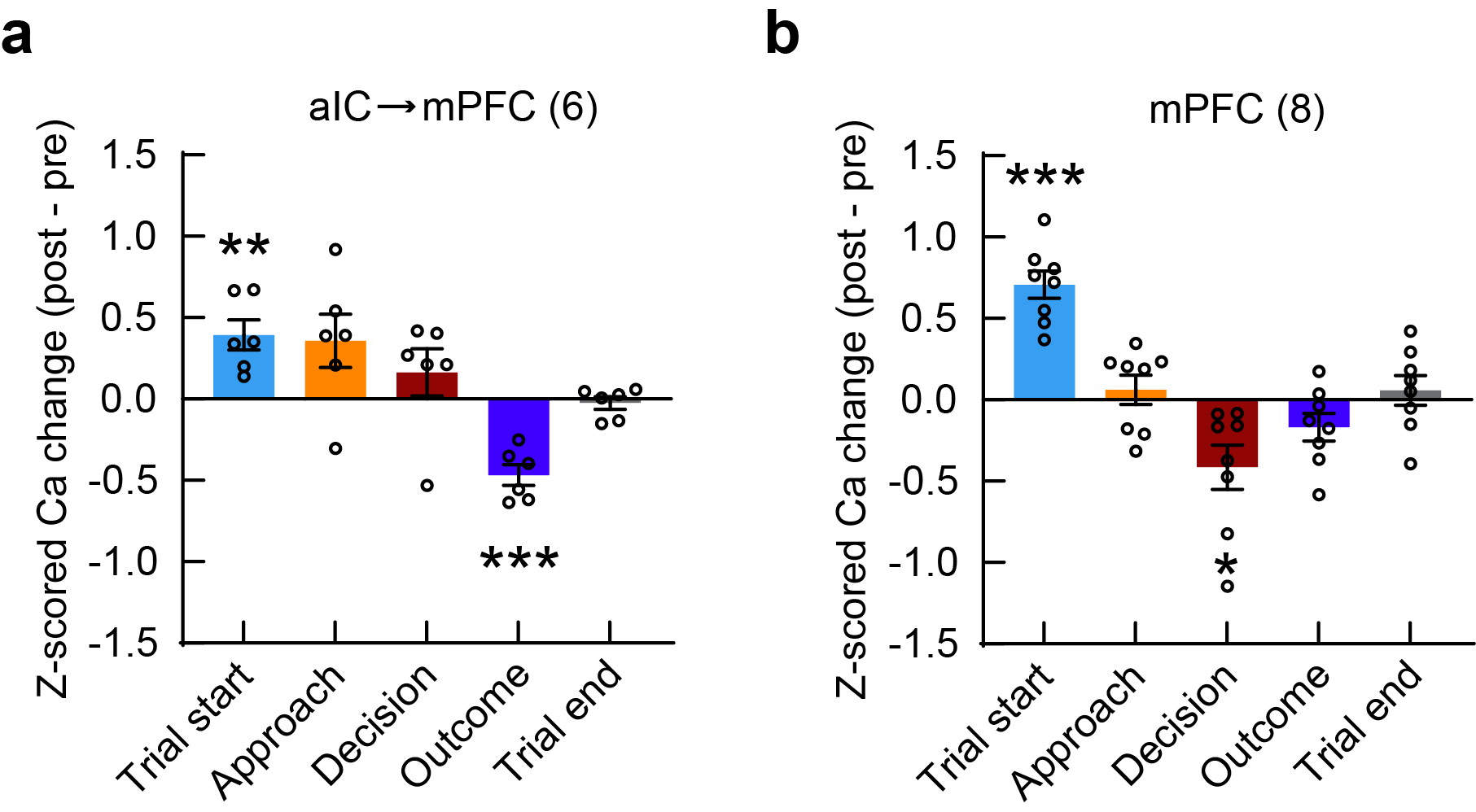

**Extended Data Figure 4. Activity of aIC→mPFC neurons and mPFC neurons during Rev measured by fiber photometry.** Ca^2+^ activity changes in aIC→mPFC neurons (**a**) and mPFC neurons (**b**) aligned with key behavioral events. One-sample *t*-test compared to zero. In **a**, Trial start: *t*(5) = 4.241, *p* = 0.0082; Approach: *t*(5) = 2.176, *p* = 0.0815; Decision: *t*(5) = 1.138, *p* = 0.3068; Outcome: *t*(5) = 7.271, *p* = 0.0008; Trial end: *t*(5) = 0.6424, *p* = 0.5489. In **b**, Trial start: *t*(7) = 8.464, *p* < 0.0001; Approach: *t*(7) = 0.6865, *p* = 0.5154; Decision: *t*(7) = 3.030, *p* = 0.0191; Outcome: *t*(7) = 2.001, *p* = 0.0855; Trial end: *t*(7) = 0.6401, *p* = 0.5425. Data are presented as mean ± s.e.m. Same mice as in Fig. 2. Sample sizes (number of mice) are given in the graphs. **p*<0.05, ***p*<0.01, ****p*<0.001.

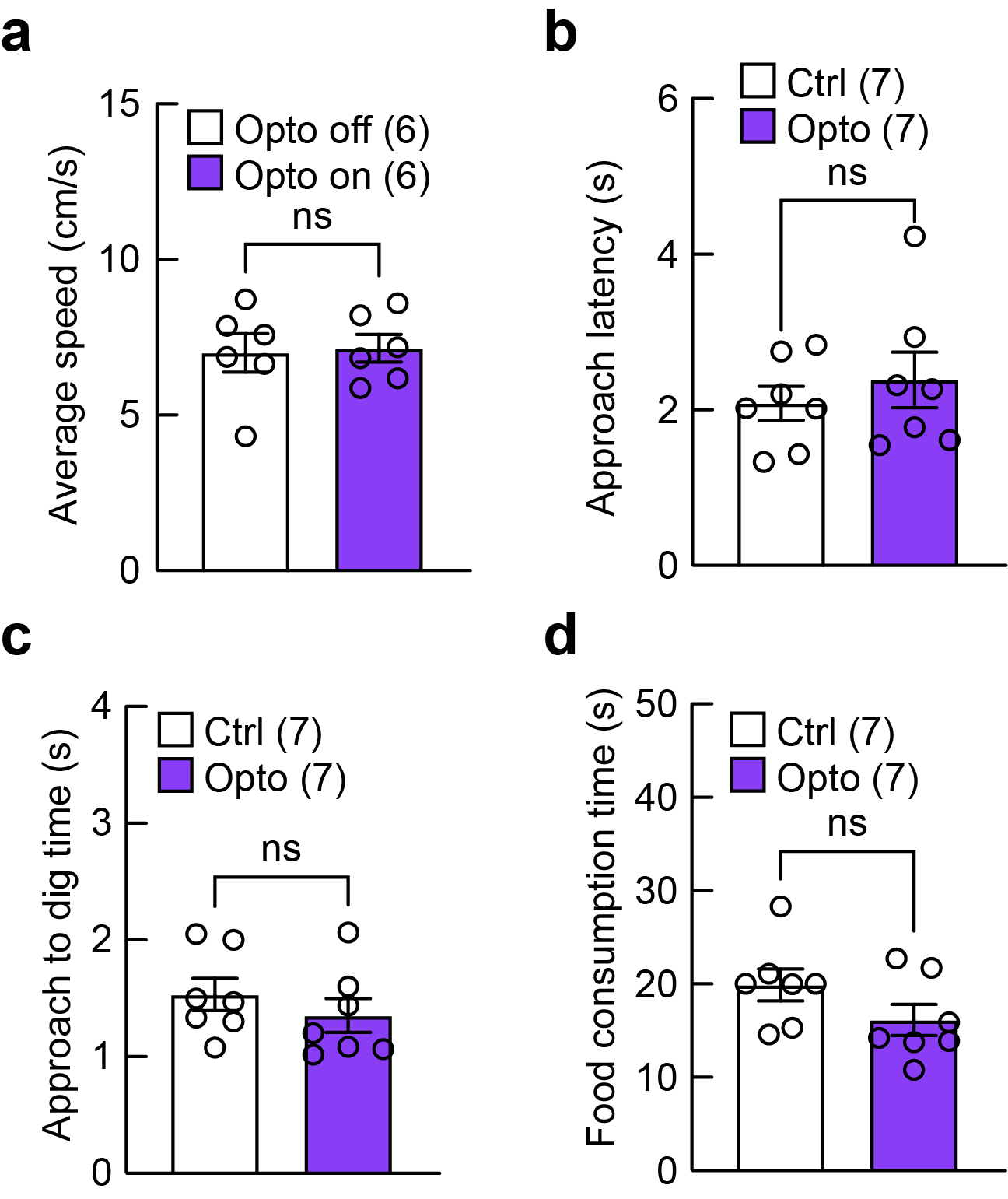

**Extended Data Figure 5. Optogenetic activation of aIC→mPFC projections did not** **alter basic motor behaviors.** Opto-stimulation did not affect locomotion (**a**), nor did it affect approach latency (**b**), approach-to-dig time (**c**), or food consumption (**d**) during EDS. Ctrl: Control; Opto: Optogenetic activation of aIC terminals in mPFC. **a,** paired t-test: *t*(5) = 0.3807, *p* = 0.7191; **b-d**, unpaired t-test: **b**, *t*(12) = 0.7142, *p* = 0.4888; **c**, *t*(12) = 0.9056, *p* = 0.3830; **d**, *t*(12) = 1.570, *p* = 0.1425. Data are presented as mean ± s.e.m. Sample sizes (number of mice) are given in the graphs. ns: not significant.

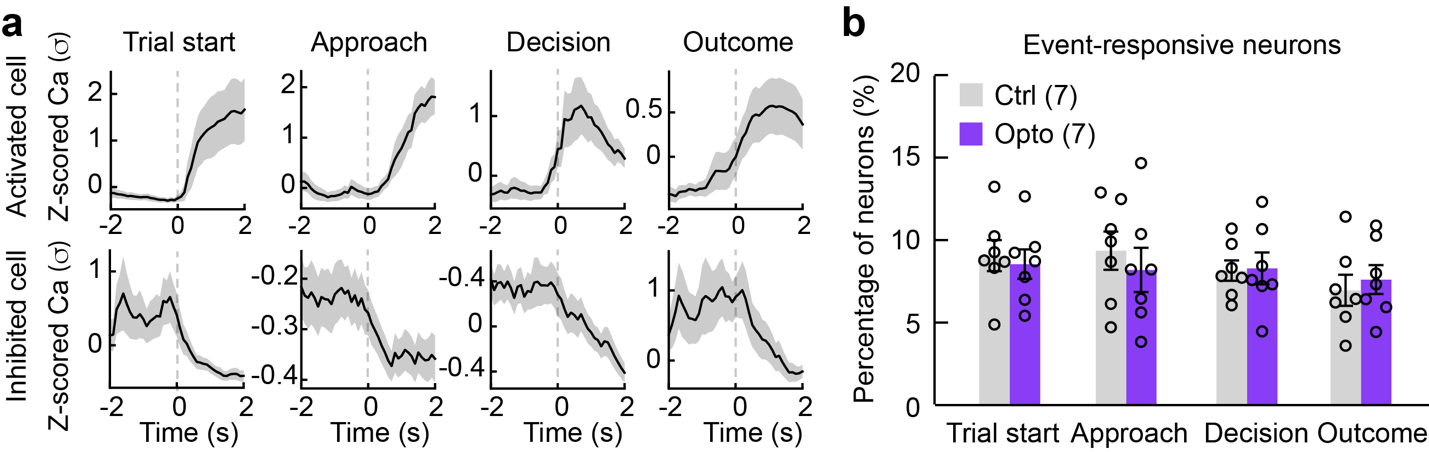

**Extended Data Figure 6. Optogenetic activation of aIC→mPFC projections did not change the proportion of mPFC neurons modulated by different behavioral events. a**, Representative Ca^2+^ traces of mPFC neurons modulated by different behavioral events in EDS. Top: activated neurons; bottom: inhibited neurons. Solid lines: mean over trials, shadows: s.e.m. **b**, Percentages of mPFC neurons selectively modulated by various behavioral events. n = 7 mice per condition. Two-way repeated measures ANOVA followed by Sidak’s multiple comparisons. Main effect for Treatment: *F*(1,12) = 0.1165, *p* = 0.7387; Behavior event: *F*(3,36) = 0.9999, *p* = 0.4040; interaction between Treatment and Behavior event: *F*(3,36) = 0.3067, *p* = 0.8204. Ctrl: Control; Opto: Pre-decision optogenetic activation of aIC terminals in mPFC. Data are presented as mean ± s.e.m. Sample sizes (number of mice) are given in the graphs.

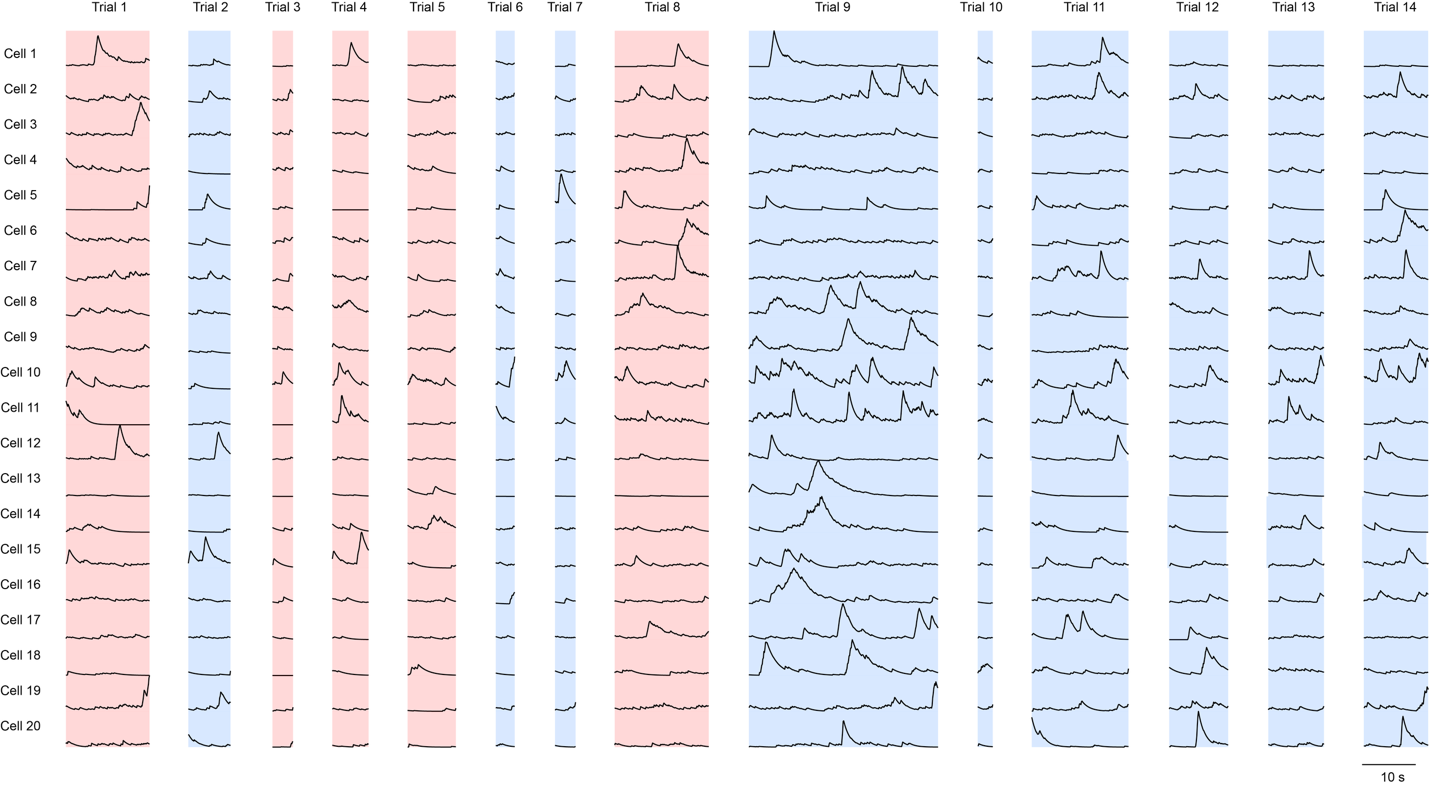
**Extended Data Figure 7. Representative Ca2+ traces of individual neurons from trial start to decision across all EDS trials.** Correct trials are shaded in blue, and incorrect trials are shaded in red.

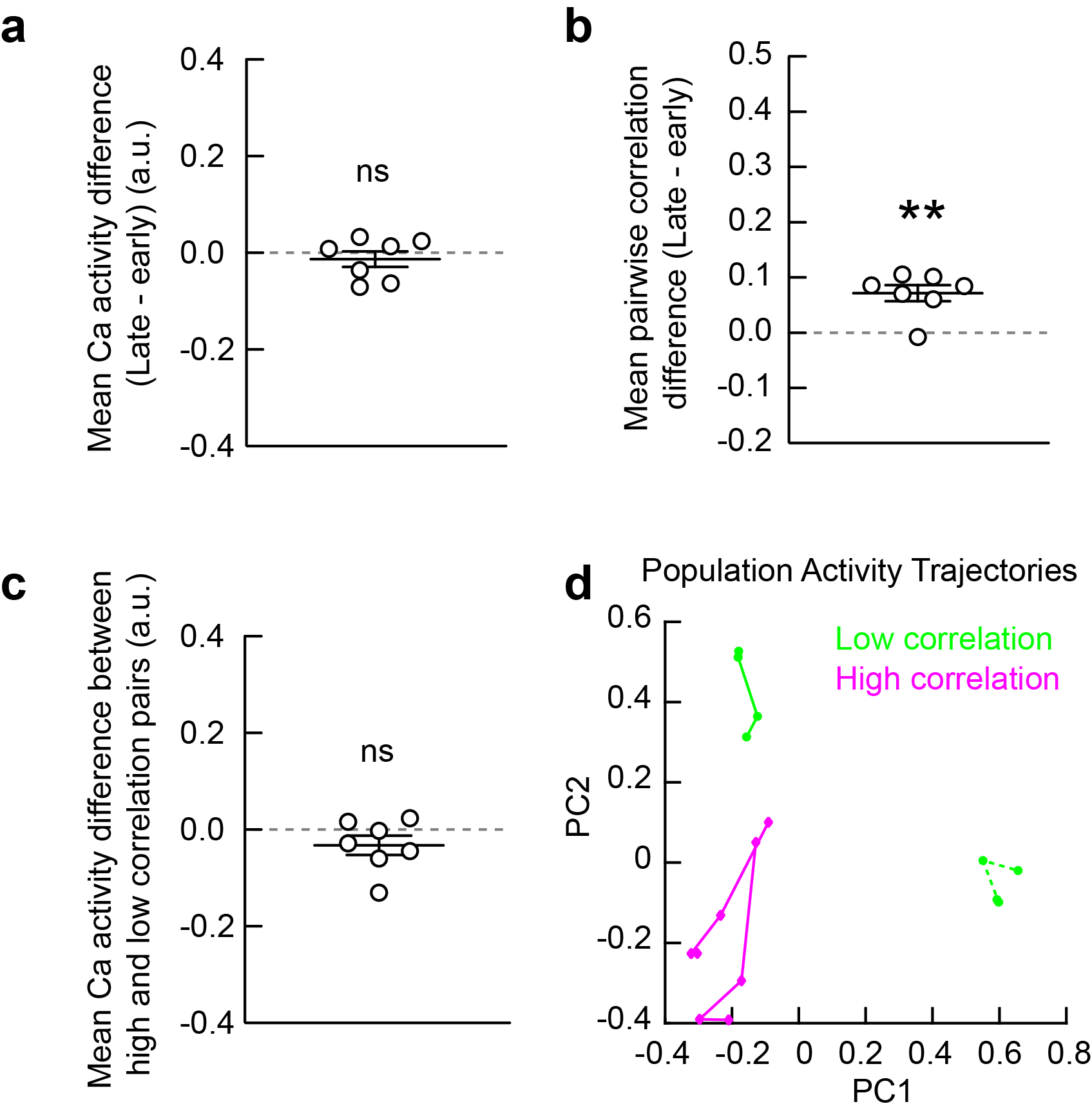

**Extended Data Figure 8. Relationship between trial-to-trial population correlation and neuronal activity in mPFC. a, b,** Early and late correct trials during EDS show comparable levels of mean population activity (**a**) but differ significantly in the average pairwise Pearson correlation (**b**). The first 6 correct trials are defined as “Early”, and the last 6 correct trials are defined as “Late”. One-sample t-test compared to 0; **a**, *t*(6) = 0.7963, *p* = 0.4562; **b**, *t*(6) = 4.932, *p* = 0.0026. **c,** High-correlation trial pairs (defined as those with Pearson correlation in the top 5% among all pairs) do not exhibit significantly higher average neural activity level than low-correlation trial pairs (defined as those with Pearson correlation in the bottom 5%). One-sample t-test compared to 0; *t*(6) = 1.603, *p* = 0.1601. **d,** Example trajectories of population activity from a single mouse, projected onto the plane spanned by the first two PCA axes. Trajectories from one pair of consecutive correct trials with high correlation (magenta) are closely aligned, while those from a low-correlation pair beginning with an incorrect trial (green) are more divergent. Each colored line represents the trajectory of a single trial, with each dot indicating the average population activity within one of the 4 pre-decision time-windows. Solid lines: correct trials; dotted lines: incorrect trials. Data in **a–c** are presented as mean ± s.e.m. n = 7 mice. ***p* < 0.01. ns: not significant. a.u.: arbitrary units.

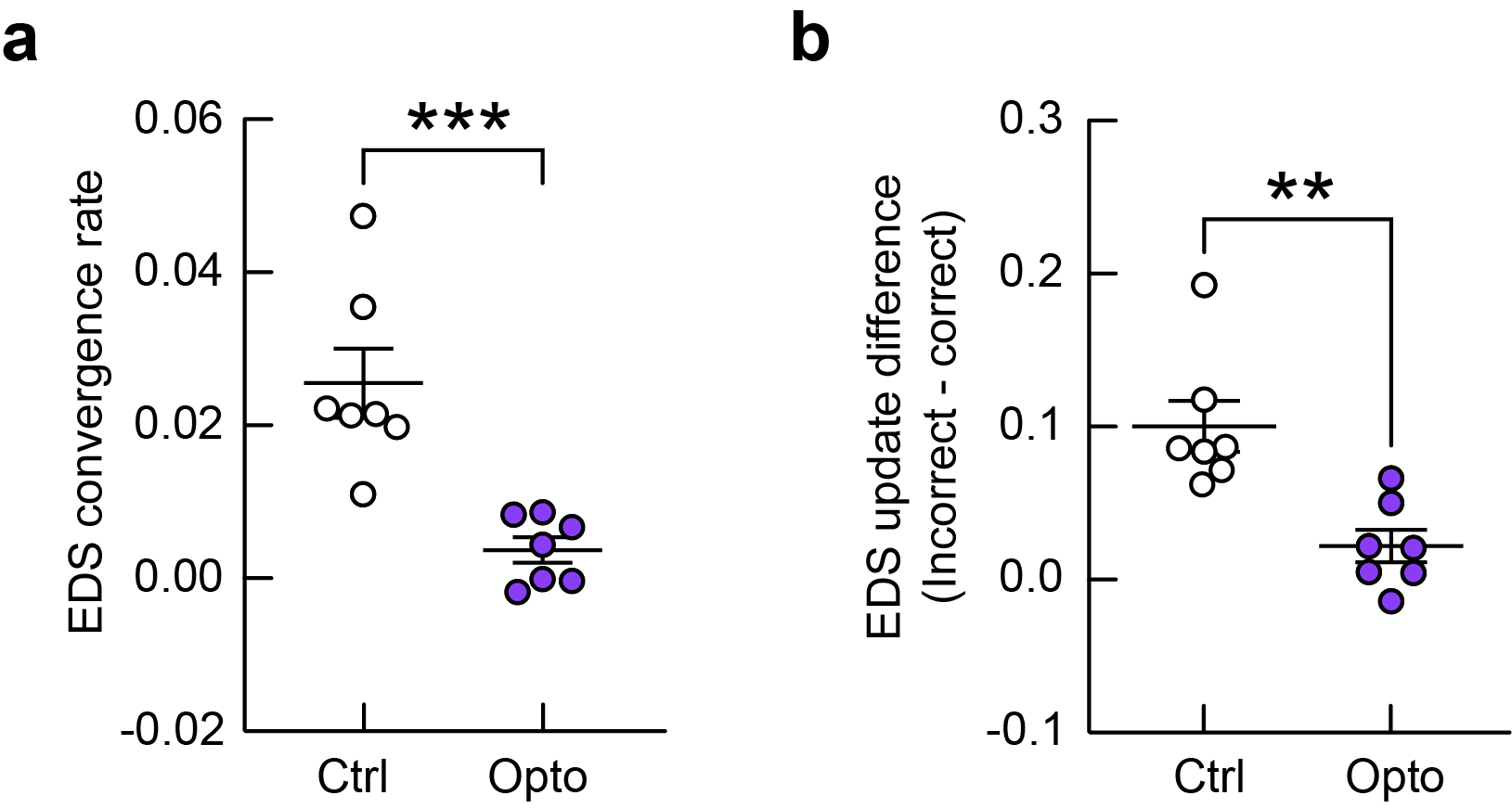

**Extended Data Figure 9. Spearman rank correlation analysis revealed population activity pattern convergence and outcome-dependent update differences consistent with those identified using Pearson correlation.** **a**, EDS convergence rate across trials in Ctrl and Opto mice. Unpaired t-test: *t*(12) = 4.517, *p* = 0.0007. **b**, EDS update difference in Ctrl and Opto mice. Unpaired t-test: *t*(12) = 3.953, *p* = 0.0019. n = 7 mice per condition. Same animals as show in Fig. 3g and 3j. Data are presented as mean ± s.e.m. ***p*<0.01, ****p* <0.001.

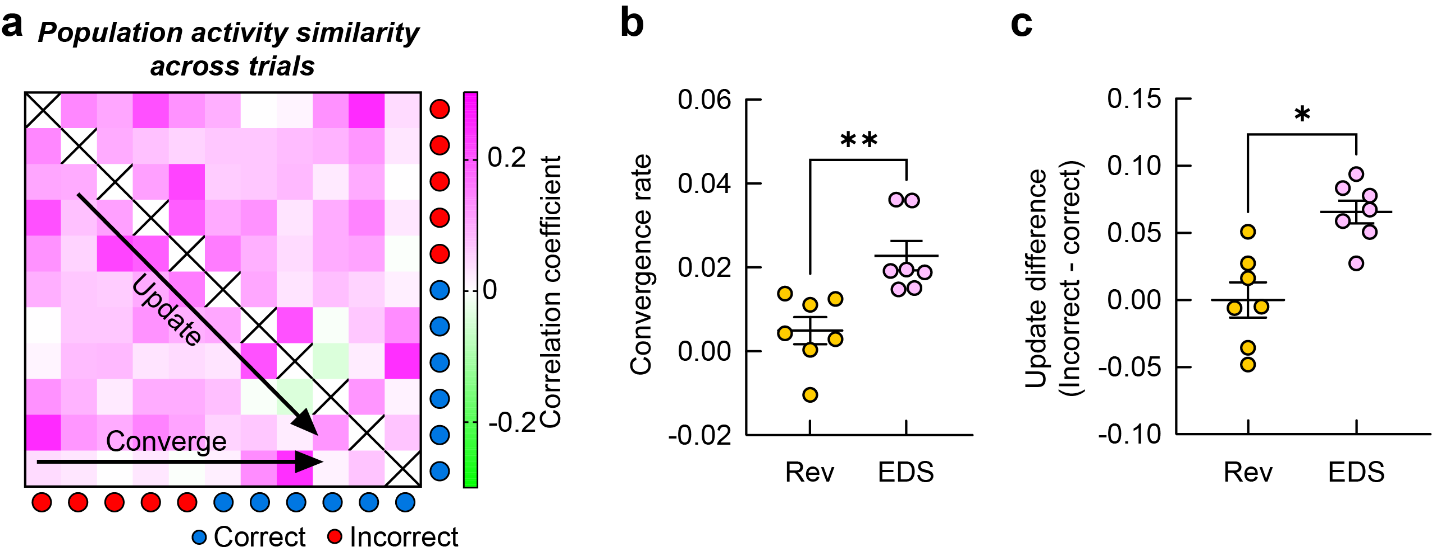

**Extended Data Figure 10. Convergence and update of mPFC neuronal population activities across Rev trials in control mice. a,** Correlation matrix showing pairwise comparisons of mPFC population activity patterns during Rev trials. **b**, Convergence rate of population activity patterns is significantly higher in EDS than in Rev. Paired t-test: *t*(6) = 5.012, *p* = 0.0024. **c**, Update difference in EDS is significantly higher than that in Rev. Paired t-test: *t*(6) = 3.220, *p* = 0.0181. n = 7 mice. EDS data are same as that in Fig. 3g and 3j. Data are presented as mean ± s.e.m. **p*<0.05, ***p*<0.01.

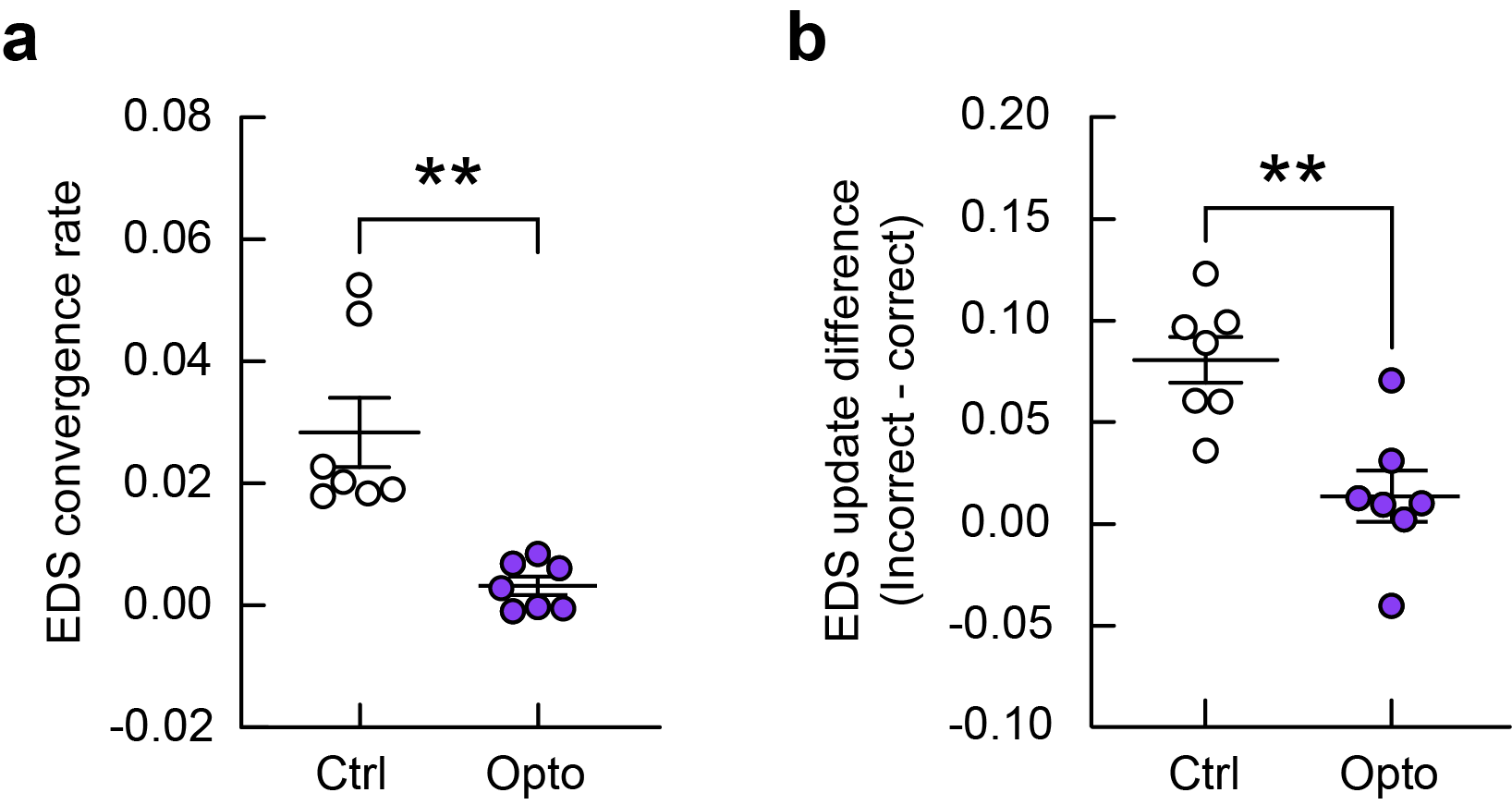

**Extended Data Figure 11. The effects of optogenetic manipulation remain evident even when neuronal activity is averaged within each pre-decision time window.** Each cell’s activity (recorded at 10 Hz) was averaged within four 2-second time windows preceding the decision-making event (post-trial start, pre-approach, post-approach, and pre-digging). **a**, Optogenetic stimulation of the aIC→mPFC projection significantly reduces the convergence rate of mPFC population activity patterns during EDS trials. Unpaired t-test: *t*(12) = 4.287, *p* = 0.0011. **b**, Optogenetic stimulation of the aIC→mPFC projection decreases the trial-outcome dependence of mPFC update. Unpaired t-test: *t*(12) = 3.979, *p* = 0.0018. n = 7 mice per condition. Same animals as shown in Fig. 3g and 3j. Data are presented as mean ± s.e.m. ***p*<0.01.

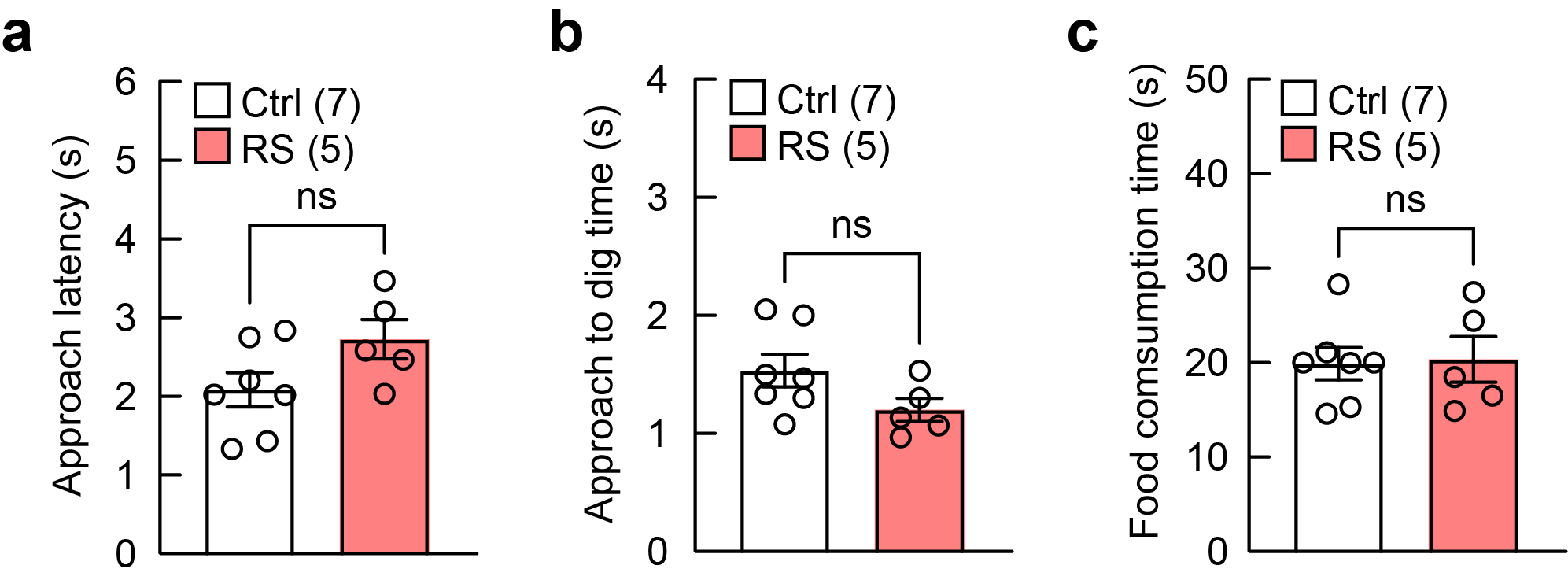

**Extended Data Figure 12. Stressed mice show normal approach latency (a), approach to dig time (b) and food consumption time (c) during EDS.** Ctrl group is the same presented in Extended Data Fig. 5b-d. Unpaired t-test: **a**, *t*(10) = 1.917, *p* = 0.0842; **b**, *t*(10) = 1.807, *p* = 0.1009; **c**, *t*(10) = 0.1592, *p* = 0.8767. Data are presented as mean ± s.e.m. Sample sizes (number of mice) are given in the graphs. ns: not significant.

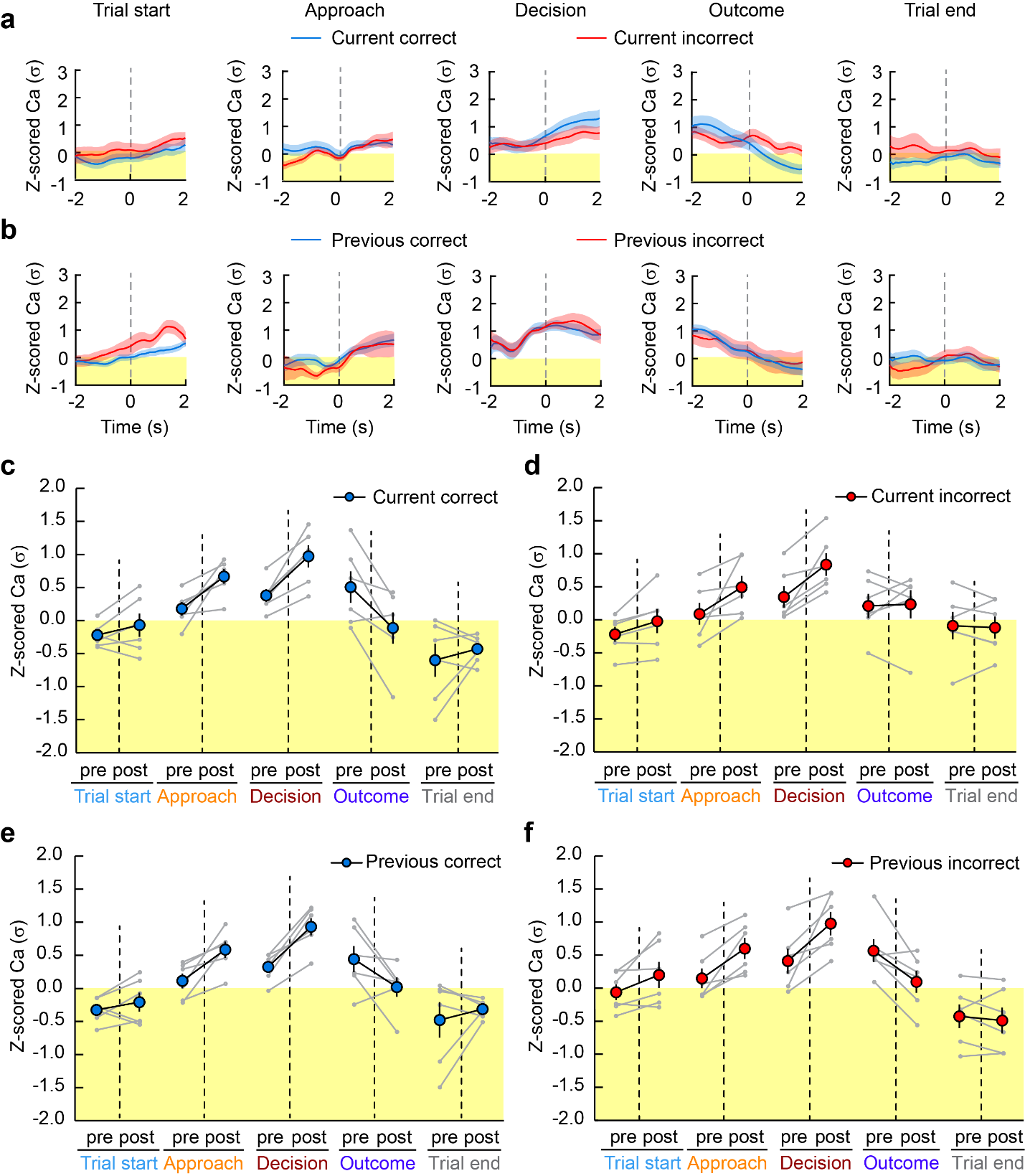

**Extended Data Figure 13. Representative Ca^2+^ traces and data points of individual animals obtained by fiber photometry in stressed mice. a,b,** Representative Ca^2+^ traces of aIC→mPFC neurons in stressed mice during different behavioral events, separated by current (**a**) or previous (**b**) trial outcome. Solid lines: mean; shadows: s.e.m.. **c-f**, Ca^2+^ activity of aIC→mPFC neurons in stressed mice, separated by current trial’s outcome (**c,d**) or previous trial’s outcome (**e,f**). Data are shown as mean ± s.e.m.
